# Repeated behavioral evolution is associated with targeted convergence of gene expression in cavity-nesting songbirds

**DOI:** 10.1101/2024.02.13.580205

**Authors:** Sara E Lipshutz, Mark S Hibbins, Alexandra B Bentz, Aaron M Buechlein, Tara A Empson, Elizabeth M George, Mark E Hauber, Doug B Rusch, Wendy M Schelsky, Quinn K. Thomas, Samuel J. Torneo, Abbigail M Turner, Sarah E Wolf, Mary J Woodruff, Matthew W Hahn, Kimberly A Rosvall

## Abstract

Uncovering the genomic bases of phenotypic adaptation is a major goal in biology, but this has been hard to achieve for complex behavioral traits. Here, we leverage the repeated, independent evolution of obligate cavity-nesting in birds to test the hypothesis that pressure to compete for a limited breeding resource has facilitated convergent evolution in behavior, hormones, and gene expression. We used an integrative approach, combining aggression assays in the field, testosterone measures, and transcriptome-wide analyses of the brain in wild-captured females and males. Our experimental design compared species pairs across five avian families, each including one obligate cavity-nesting species and a related species with a more flexible nest strategy. We find behavioral convergence, with higher levels of territorial aggression in obligate cavity-nesters, particularly among females. Across species, levels of testosterone in circulation were not associated with nest strategy, nor aggression. Phylogenetic analyses of individual genes and co-regulated gene networks revealed more shared patterns of brain gene expression than expected by drift, but the scope of convergent gene expression evolution was limited to a small percent of the genome. When comparing our results to other studies that did not use phylogenetic methods, we suggest that accounting for shared evolutionary history may reduce the number of genes inferred as convergently evolving. Altogether, we find that behavioral convergence in response to shared ecological pressures is associated with largely independent gene expression evolution across different avian families, punctuated by a narrow set of convergently evolving genes.

## Main

Biologists have long been fascinated by phenotypic convergence as a window into the predictability of evolution (1-3). However, our understanding of the molecular mechanisms of behavioral convergence lags behind that of morphological or physiological traits (4-6). Some studies of behavioral convergence find that evolution repeatedly uses the same (or similar) changes in particular genes or pathways (7-9). Alternatively, the building blocks of behavioral convergence may be independent, with lineage-specific mechanisms across replicated evolutionary events (10, 11). Understanding the relative contributions of these two processes requires a comparative approach that embraces the likely polygenic nature of complex behaviors (12-16). However, efforts to connect behavioral evolution to transcriptomics rarely use phylogenetic methods (6), despite the potential for changes in gene expression to shape phenotypic diversification. Likewise, genetic drift may shape gene expression independently of adaptation, and phylogenetic methods are necessary to disentangle these processes.

Territorial aggression is a widespread behavioral trait that is well-suited for evaluating these processes. In vertebrates, aggression is mediated by diverse neuroendocrine mechanisms (17), including the hormone testosterone (18) and its metabolite 17β-estradiol, both of which can act on sex steroid receptors in the brain to promote aggression (19, 20). Aggression is also linked to brain metabolic pathways (7) and G-protein coupled receptor signaling of dopamine, serotonin, and glutamate (21, 22). Many of these candidates are shared across distantly related species (7, 21), demonstrating the potential for conserved or repeatedly evolved mechanisms in the evolution of territorial aggression. However, there is also interspecific variation in the abundance and distribution of these endocrine-molecular building blocks (23, 24), indicating that mechanisms of aggression can diverge over time.

Mechanisms of aggression can also differ within a species, including between males and females (25), both of whom exhibit territorial aggression in many species (26, 27). This provides a unique opportunity to include both sexes in analyses of behavioral evolution. After all, females and males share the majority of their genome, yet sex-specific selective pressures can alter gene expression (28) and hormone secretion (29), which together may shape aggression or the mechanisms that promote the expression of aggression, the latter of which is still remarkably understudied in females (30, 31). Critically, in both sexes, aggression shapes access to resources and mates, and the degree of such competition varies among species (26, 27).

Competition for breeding territories is thought to be especially intense for obligate secondary cavity-nesting birds, which must secure an excavated space inside a tree or other substrate. They need a cavity to breed, they cannot excavate one themselves, and physical conflicts over cavities can lead to serious injury or death (32-34). For many cavity-nesting species, both females and males compete to acquire and defend nesting territories (35-39), and our earlier case study showed elevated aggression in two species of obligate cavity-nesters, though without quantitative phylogenetic methods (40). There have been multiple evolutionary transitions in and out of secondary cavity-nesting across passerines (41, 42), providing a strong foundation to evaluate the degree of mechanistic convergence in behavioral evolution.

Here, we test the hypothesis of evolutionary convergence using a phylogenetic approach that integrates behavioral, hormonal, and neurogenomic data. We compare species pairs from five avian families - Hirundinidae (Swallows), Parulidae (Woodwarblers), Passeridae (Sparrows), Turdidae (Thrushes), and Troglodytidae (Wrens) - which each represent independent origins of obligate secondary cavity-nesting (Figure 1A; further details in SI §1) (42). Each pair includes one obligate secondary cavity-nesting species, which regularly nests in artificial nest boxes, and one species with a more flexible nest strategy, either open cup-nesting or facultative cavity-nesting. Each species pair is similar in other aspects of its ecology and life history (e.g. foraging ecology, degree of biparental care). Across species, we compared territorial aggression, testosterone in circulation, and brain gene expression. By applying quantitative phylogenetic approaches across these datasets, we evaluate molecular convergence against a null hypothesis of shared evolutionary history and genetic drift.

**Figure 1:**
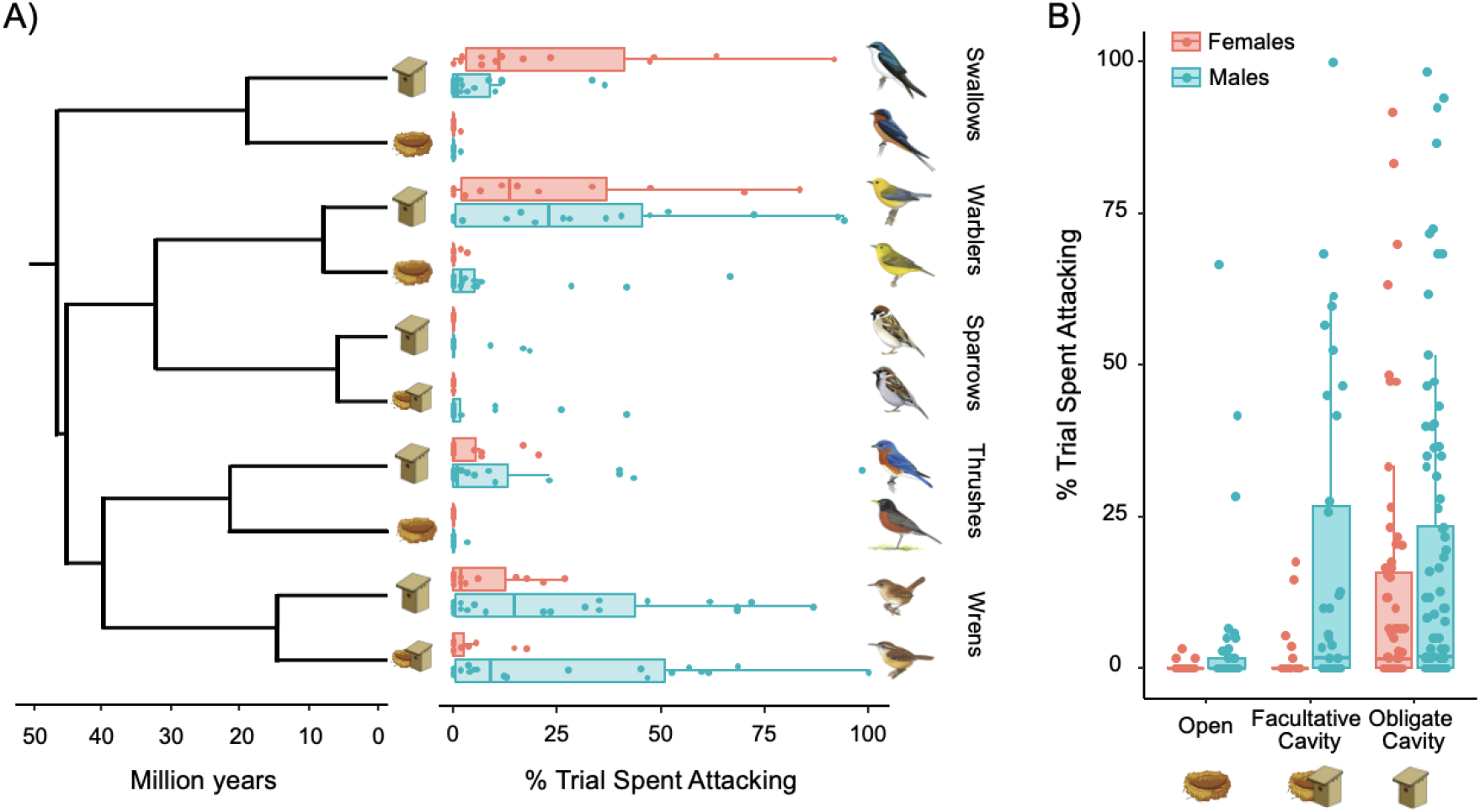
**(A)** *Left*: Consensus phylogeny of 10 species from five families, which diverged from a common ancestor ∼44 mya (36-50 mya). Species pairs diverged ∼9-20 mya (43). Nestboxes indicate obligate cavity-nesters, nests represent open nesters, and both symbols together represent facultative cavity-nesters. *Right*: Sex- and species-level aggression towards a conspecific decoy, measured by the proportion of 5-sec intervals that contained physical contact during a 5-min aggression assay. Species listed in descending order: Swallows: tree swallows (*Tachycineta bicolor*), barn swallows (*Hirundo rustica*), Woodwarblers: prothonotary warblers (*Protonotaria citrea*), yellow warblers (*Setophaga petechia*), Sparrows: Eurasian tree sparrows (*Passer montanus*), house sparrows (*Passer domesticus*), Thrushes: Eastern bluebirds (*Sialia sialis*), American robins (*Turdus migratorius*), and Wrens: house wrens (*Troglodytes aedon*), Carolina wrens (*Thryothorus ludovicianus*). **(B)** Aggression, grouped by nest strategy. Each point is one assay on a unique free-living individual; box plots convey interquartile range. Obligate cavity-nesting females were significantly more aggressive than females with more flexible nest strategies. Illustrations reproduced with permission from Lynx Edicions.

## Results

### Behavioral convergence among obligate cavity-nesters

Our study occurred during territorial establishment at the beginning of the breeding season, when birds are defending territories and acquiring mates, but before egg laying. We assayed aggression in 304 free-living female and male birds in Indiana, Illinois, or Kentucky, USA (SI §1, Table 1). Each aggression assay lasted 5 minutes and included a conspecific, taxidermic decoy of either female or male sex, coupled with playback of female or male vocalizations that occur during natural aggressive interactions (details in SI §2). We measured a variety of aggressive behaviors and focused on physical attacks (i.e. contact with the decoy). To evaluate the role of nest strategy as a driver of territorial aggression, we used phylogenetic generalized linear mixed models (PGLMMs). Briefly, these models correct for covariation in traits among species due to common ancestry by incorporating phylogenetic relatedness as a random effect. Along with sex of the territory holder and sex of the decoy, we included nest strategy as a fixed effect with two levels: obligate cavity-nesting vs. more flexible strategies (including both facultative cavity- and open-nesting). We also ran three-level models differentiating obligate cavity, facultative cavity, and open nest strategies (SI §9-11).

**Table 1.**
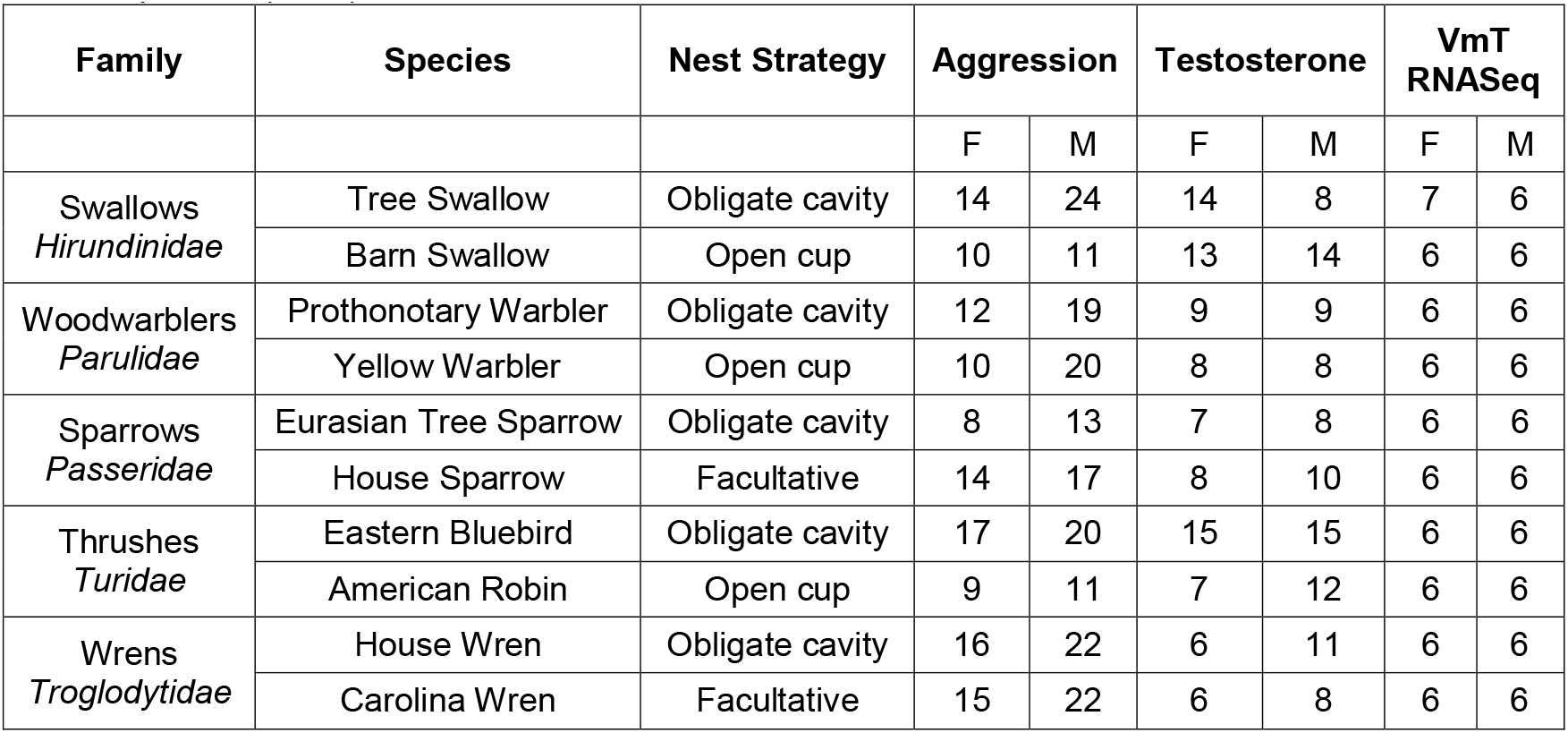
Sample sizes for behavior, testosterone, and gene expression in the ventromedial telencephalon (VmT).

| Family | Species | Nest Strategy | Aggression |  | Testosterone |  | VmT RNASeq |  |
| --- | --- | --- | --- | --- | --- | --- | --- | --- |
|  |  |  | F | M | F | M | F | M |
| Swallows<br>*Hirundinidae* | Tree Swallow | Obligate cavity | 14 | 24 | 14 | 8 | 7 | 6 |
|  | Barn Swallow | Open cup | 10 | 11 | 13 | 14 | 6 | 6 |
| Woodwarblers<br>*Parulidae* | Prothonotary Warbler | Obligate cavity | 12 | 19 | 9 | 9 | 6 | 6 |
|  | Yellow Warbler | Open cup | 10 | 20 | 8 | 8 | 6 | 6 |
| Sparrows<br>*Passeridae* | Eurasian Tree Sparrow | Obligate cavity | 8 | 13 | 7 | 8 | 6 | 6 |
|  | House Sparrow | Facultative | 14 | 17 | 8 | 10 | 6 | 6 |
| Thrushes<br>*Turdidae* | Eastern Bluebird | Obligate cavity | 17 | 20 | 15 | 15 | 6 | 6 |
|  | American Robin | Open cup | 9 | 11 | 7 | 12 | 6 | 6 |
| Wrens<br>*Troglodytidae* | House Wren | Obligate cavity | 16 | 22 | 6 | 11 | 6 | 6 |
|  | Carolina Wren | Facultative | 15 | 22 | 6 | 8 | 6 | 6 |

We found significant effects of nest strategy, sex of the territory holder, sex of the decoy, and their interaction on aggression. Obligate cavity-nesters spent more time attacking the decoy, compared to species with more flexible nest strategies (PGLMM: p = 0.0011; Figure 1A; Table S1). This difference in aggression was larger between obligate cavity-nesters and open nesters relative to facultative cavity-nesters (Figure 1A). Males attacked marginally more than females (PGLMM: p = 0.058). Notably, sex interacted with nest strategy (PGLMM: p = 0.0064), a pattern driven by elevated levels of territorial aggression in obligate cavity-nesting females (Figure 1B). We also found a significant interaction between sex of the territory holder and sex of the decoy (PGLMM: p = 0.011), such that male territory holders were more aggressive towards male decoys, whereas female territory holders attacked female and male decoys with similar levels of aggression (Figure S1; Table S1).

Additionally, we measured the average distance from the focal individual to the decoy, to confirm each subject was present and engaged during a trial, regardless of whether they were aggressively attacking. Distance from the decoy was not related to nest strategy (PGLMM: p = 0.22), sex (p = 0.70), nor their interaction (p = 0.39) (Table S2; Figure S2). This result indicates that decoy placement was a salient stimulus, in that all species were in audio-visual proximity to the simulated intruder (average distance = 5.0m), but obligate cavity-nesters spent more time attacking the decoy compared to their close relatives with more flexible nesting strategies (Figure 1, Table S1).

### Testosterone levels are not associated with nest strategy, nor aggression, across species

Next, we sought to measure levels of testosterone and gene expression as potential physiological drivers of behavioral convergence. We focused our analyses on constitutive hormonal and neurogenomic states that are representative of unprovoked, free-living animals (n = 196 hormone samples, n = 121 brain gene expression samples; Table 1). We followed the assumption that social elevation of testosterone peaks around 30-45 min after HPG axis activation (44, 45), and that socially responsive genes likewise show peak transcriptional responses 30-60 minutes after a stimulus (46). To maximize our number of aggression assays while meeting these assumptions, our sample collection took three approaches: a) passive collection, in which we set up a mist-net or trap in the target territory and waited to capture the focal individual without any stimulation (testosterone: n = 108, brain: n = 51), b) immediate collection, in which we sampled individuals immediately after a short aggression assay (testosterone: n = 55, brain: n = 43), and c) delayed collection in which we sampled individuals several days after a short aggression assay (testosterone: n = 33, brain: n = 27). We found no differences in testosterone among immediate, passive, or delayed sample collection approaches (ANOVA: F_2, 184_ = 0.011, p = 0.90; Figure S3A), and therefore combined these samples.

We previously identified a positive correlation between territorial aggression and testosterone within female tree swallows (S = 2, p = 0.0028, rho = 0.96) (40), which are included in this study, but we did not find this relationship in male tree swallows. To explore whether similar patterns apply to other species sampled here, we conducted Pearson’s correlations across species for each sex, and Spearman’s correlations within each family for each sex, given the reduced sample size. We did not find any relationship between testosterone and aggression in Eastern bluebirds, house wrens, Carolina wrens, prothonotary warblers, nor yellow warblers (p > 0.13), and we did not have enough data on testosterone and aggression from the same individuals for the other species. Considered collectively for all samples, we did not find a significant relationship between territorial aggression and testosterone for females (t = 1.87, df = 25, p = 0.073, r = 0.35), nor for males (t = 1.17, df = 35, p = 0.25, r = 0.19). Testosterone was not related to nest strategy (PGLMM: p = 0.80) nor the interaction between nest strategy and sex (PGLMM: p = 0.85; Figure S4, Table S3). As expected, we found that males had significantly higher levels of testosterone in circulation than females (PGLMM: p < 0.0001).

### Neurogenomic mechanisms of behavioral convergence

Finally, we examined convergent evolution in brain gene expression. Using RNA-seq, we measured mRNA abundance for 10,672 orthologous genes expressed in all 10 focal species in the ventromedial telencephalon (Table 1, Figure S5); this region contains core nodes of the vertebrate social behavior network, which regulates behaviors including aggression (47). The number of differentially expressed genes generally increased with divergence time between species pairs within each family (p = 0.08, R^2^ = 0.83; SI §5, Figure S8, Table S4), underscoring the need for phylogenetic methods. Using these expression data, we took three analytical approaches to understand the mechanisms underlying convergent behavioral evolution.

### Gene expression is highly concordant between pairs of families, but involves different sets of genes across the phylogeny

We used the Rank Rank Hypergeometric Overlap (RRHO) approach to compare the magnitude and direction of gene expression differences among species pairs. If the same genes are repeatedly targeted by selection for the obligate cavity-nesting strategy, these genes should be differentially expressed in the same direction for cavity-nesters across multiple family comparisons. As a null hypothesis, we also generated permuted differential expression datasets by randomly sampling the log2FoldChange and associated p-values from the observed values with replacement for each gene. This was done independently for each family. The permuted datasets were then given to the same RRHO pipeline used for the empirical datasets.

In contrast to the largely random distributions of overlap from the randomly permuted dataset (Figure S9), RRHO for our empirical dataset revealed significantly concordant patterns of gene expression (all p-values < 0.01, permutation test, Figure 2, Table S5). Genes that were more highly expressed in the obligate cavity-nester of one species pair were also more highly expressed in the obligate cavity-nester of another species pair, and vice-versa for genes with lower expression (Data S4), evidenced by clustering in the upper-right and lower-left quadrants of the RRHO heatmaps (Figure 2C). This concordant pattern applied to an average of 1390 genes per family comparison (Figure S12), amounting to ∼13.0% of genes studied, suggesting a large number of genes evolving in a similar direction of expression within each family. In comparison, the randomly permuted dataset had an average of 562 concordant genes per family comparison (∼5.3% of total genes). The empirical dataset had more than twice as many concordant genes as expected by random chance from the permuted dataset. Therefore, our empirical RRHO data for each family comparison are overwhelmingly concordant, beyond the null expectation.

**Figure 2:**
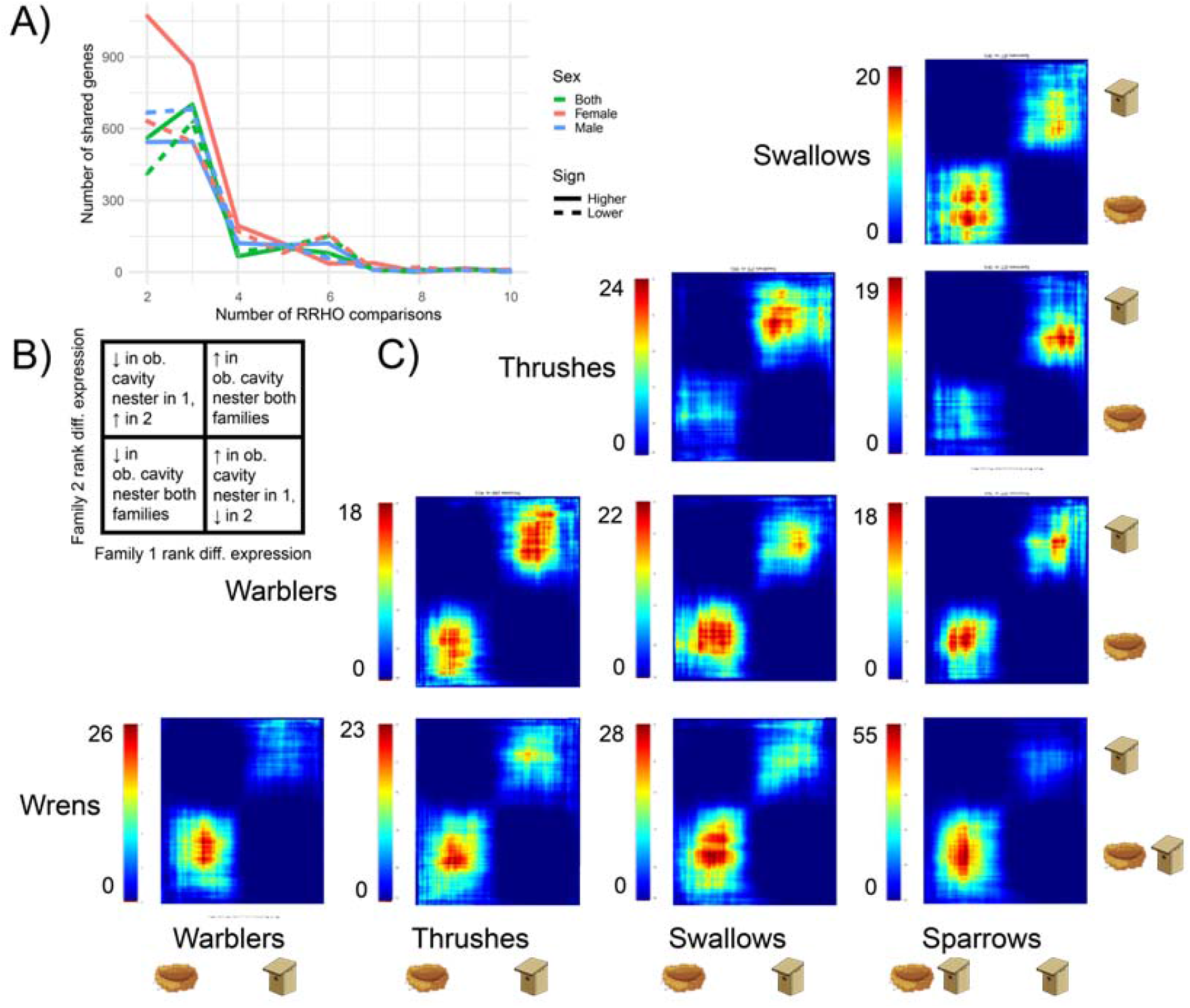
Concordance in differential expression based on rank-rank hypergeometric overlap (RRHO). Heatmap colors reflect adjusted hypergeometric -log(p-value). Larger values indicate more concordance in expression between families. Nestboxes indicate obligate cavity-nesters, nests represent open nesters, and both together represent facultative cavity nesters. **(A)** Number of concordantly expressed genes shared across family comparisons (i.e. across heatmaps) for males (blue), females (pink) and both sexes combined (green). Solid lines indicate higher expression in obligate cavity-nesters, dashed lines indicate lower expression **(B)** Key for interpreting individual heatmaps. Each quadrant of the key corresponds to a quadrant of an individual heatmap, shown in **(C)** Each pixel within the heatmaps contains two sets of approximately 100 genes being compared between the two families; the color legend indicates the log p-value of the overlap between these gene sets, with a higher value indicating stronger overlap.

This high level of concordance in gene expression did not persist across multiple family comparisons, indicating a lack of convergence in the identity of genes differentially expressed between species pairs. The number of shared genes decreased rapidly with increasing inclusion of additional family comparisons (Figure 2A). Specifically, about 1,000 genes (∼10% of orthologs) displayed concordant expression patterns among two or three families, and 497 genes (∼5%) were shared by at least 5 out of 10 family comparisons at a time (Figure 2A).

Relative to the empirical data, the randomly permuted data had more shared genes across 2 family comparisons, and fewer shared genes across 5+ family comparisons (Figure S9). Gene Ontology (GO) analyses of the 497 shared genes (Data S4, S5) revealed no significant functional enrichment. This pattern indicates that – despite a significant degree of concordance between pairs of species – concordantly expressed genes are not repeatedly recruited from any particular set of known biological processes. However, we must recognize that GO databases are biased towards model organisms, and a lack of GO enrichment does not necessarily indicate an absence of a shared biological function.

Across all ten family comparisons, only 11 genes were shared, meaning that 0.1% of 10,672 orthologs exhibited complete concordance across five independent origins of obligate cavity-nesting (Table S6). This number was still larger than the genes shared across families using the randomly permuted dataset, which was only 1 gene. Some of these 11 globally concordant genes have established connections with behavior or brain function (48). For instance, *EMC3, NPTX1*, and RGS19 are involved in neurotransmitter reception, *ATP6V0E1* and *DDX56* have ATPase activity, and most of the completely concordant genes have established connections with Alzheimer’s (*ARRDC4, CTNND1, NPTX1, WIPF2*), addiction (*RGS19*) or other neurological disorders (*AFAP1L1, PQBP1*) (48). s genes could be functionally validated in relation to cavity nesting or aggression in the future. Overall, these results from RRHO indicate a high degree of independent gene expression evolution in association with the convergent evolution of obligate cavity nesting.

We repeated these RRHO analyses with females and males separately, because sex-specific selection pressures could generate concordance that would be masked by analyzing both sexes together. We found similarly high concordance in these sex-specific analyses, in that significant gene overlap was concentrated primarily in the upper-right and lower-left quadrants. However, the degree (indicated by the scale of the heatmap) and direction (concentration of significant overlap in the upper-right vs. lower-left) of concordance differed between sexes (Figure 2A, Figure S10, S11). Males had more cases of higher expression in the obligate cavity nester of both families (upper-right), whereas females had more cases of lower expression (lower-left).

### Nest strategy and aggression are associated with convergently evolving genes

Next, we used PGLMMs to model expression of all ∼10k orthologs as a function of nest strategy, sex, aggression, sex-by-nest-strategy interactions, and aggression-by-nest-strategy interactions. To identify convergent expression evolution, these phylogenetic models explicitly account for shared expression due to evolutionary history. We expect some degree of convergence in expression evolution due to random chance (i.e. genetic drift along the phylogeny), so we employed a false discovery rate (FDR) correction to test this null hypothesis. Many genes showed some relationship with obligate cavity nesting, aggression, or a relevant interaction, supporting a hypothesis of convergent expression evolution. Specifically, there were 234 genes associated with obligate cavity-nesting, 79 genes associated with aggression, and five genes overlapped among these. Of these five genes (*RNASEH2B, ERN1, UGGT2, TAF1B*, and *HIGD1A*), the former four relate to protein or RNA processing, the latter four relate to stress responses, and all have connections with neurodegenerative disorders (48-52). An additional 62 genes exhibited an interaction between nest strategy and aggression. 76 genes exhibited an interaction between sex and obligate cavity-nesting, with approximately equal numbers of genes that had a stronger association in females vs. males (Table S7, Data S6). Sex had the largest influence on expression variation and was associated with 510 genes (Table S7), the majority of which had higher expression in males than females. Genes with sex-biased expression were significantly enriched for cellular metabolic process.

Because some nest-strategy and aggression-associated patterns may be driven by expression in one or two species, we developed additional criteria to pinpoint a robust set of convergently evolving genes. We retained only those that were (i) significantly different in expression (by t-test) between species in at least 3 out of 5 family comparisons; and (ii) different in a consistent direction (i.e. all higher or all lower expression) with respect to nest strategy in all families with a significant difference. 168 of 234 genes met these criteria of convergence for nest strategy (Data S8). Further restricting our analyses to all 5 family comparisons, we identified 40 genes (0.4% of orthologues) associated with nest strategy. Several convergent genes associated with nest strategy have some connection to ATP and mitochondrial function (*ATP1B1, PITRM1, SLC25A24*), as well as behavior or psychiatric risk (*TRMT1L, NT5C2, BCHE*) (48).

We also used PGLMMs to examine the expression of a subset of candidate genes related to testosterone and steroid hormone signaling in our transcriptome dataset, but we did not find a relationship with obligate cavity nesting (SI §9).

### Nest strategy and female aggression are associated with convergent gene networks

Complex phenotypes are often regulated by subtle but coordinated changes in gene networks, which may also be targets of selection (53), and approaches that analyze genes individually may miss important emergent properties. Therefore, we used weighted gene co-expression network analyses (WGCNA) to estimate co-expression across all 10 species (SI §8), yielding 93 networks of correlated genes (Data S10). Using each network’s eigengene (akin to a principal component of expression), we ran PGLMM with fixed effects of nest strategy, aggression, sex, and their interactions. After accounting for phylogeny, and with FDR correction, two networks showed significant associations that are unlikely to occur by drift alone. Expression in the red network was higher in males than in females (Figure S13a), and 147 of 193 genes (76%) were on the zebra finch Z chromosome. Expression in the tan4 network was associated with nest strategy (Data S9, Table S8), with lower eigenvalues in each of the five independent origins of obligate cavity-nesting (Figure 3A). Though not significantly enriched for any GO terms (Table S8), several genes in the tan4 network related to mitochondrial function and energy metabolism, cognition, and stress response (Figure 3C).

**Figure 3.**
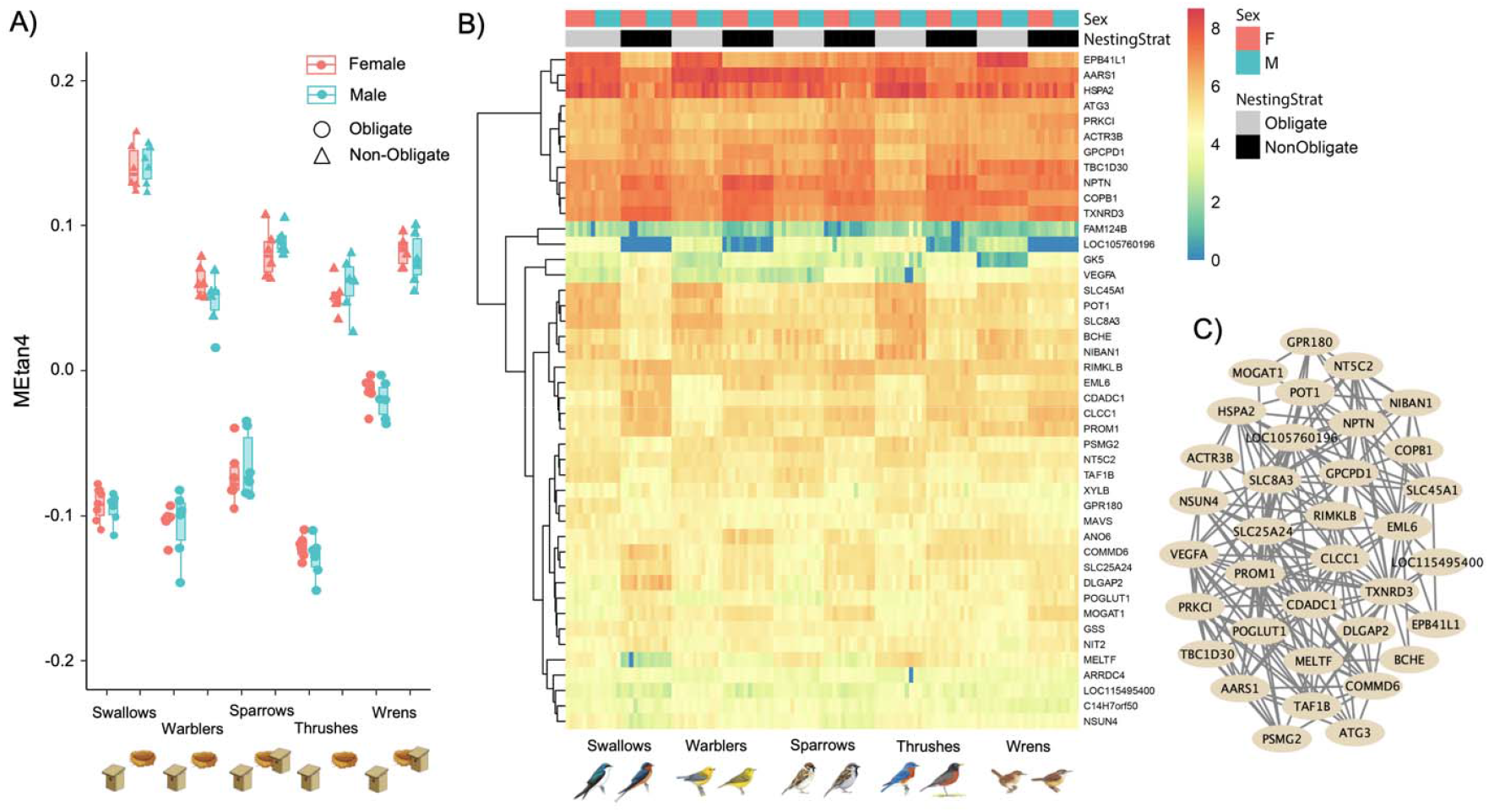
**(A)** Weighted gene co-expression network analysis (WGCNA) eigengene values for the tan4 network, which included 44 genes whose expression was significantly associated with obligate cavity-nesting relative to non-obligate nesting strategies. Individual points represent the eigengene value for a sampled individual, colored by sex. Nestboxes indicate obligate cavity-nesters, nests represent open nesters, and both together represent facultative cavity nesters. **(B)** Heatmap of log-scaled genes in tan4 network. Columns represent individuals, grouped by sex and nest strategy. Color scale indicates log of gene expression. **(C)** Network depicts genes with network membership |>| 0.6. Illustrations reproduced with permission from Lynx Edicions. Because sexes have the potential to differ in mechanisms of behavior, we repeated this analysis for each sex separately (SI §10). In this sex-specific analysis, we identified two female-specific gene networks correlated with species-level aggression – pink4 and sienna4 (Pearson’s correlation: r |>| 0.78, p < 0.0001, Figure 4, Table S9). These networks contained genes with established connections to aggression, including DRD3, a dopamine receptor in the pink4 network, and GRIA2, a glutamate receptor in the sienna4 network. Finding aggression-associated networks specific to females further suggests that there are multiple gene regulatory routes to aggression, including those that may differ between sexes.

**Figure 4.**
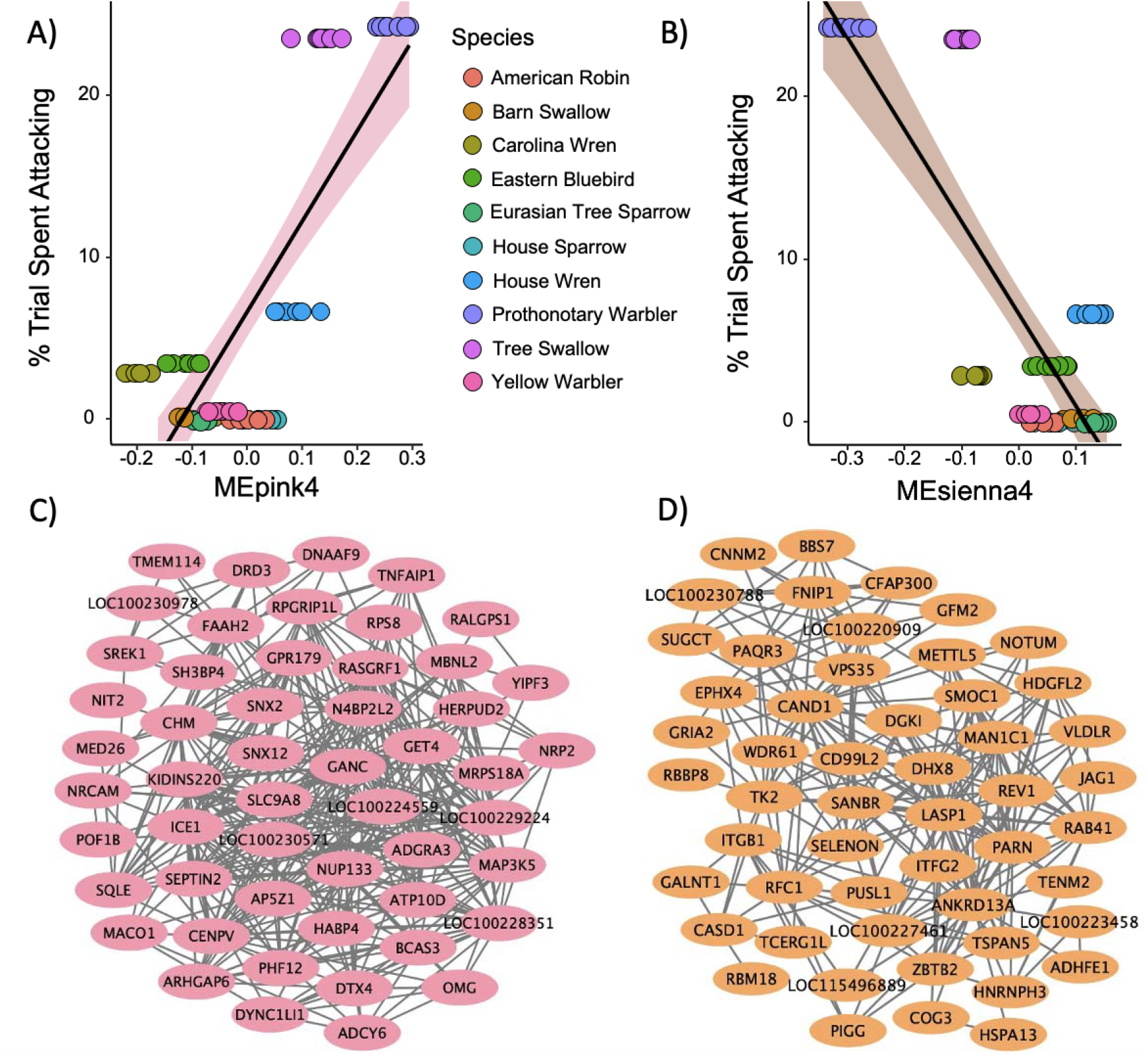
For females only, WGCNA network eigengenes for the **(A)** pink4 network and **(B)** sienna network were associated with average, species-level aggression towards a conspecific decoy, measured by the proportion of 5-sec intervals that contained physical contact during a 5-min aggression assay. Networks for **(C)** pink4 and **(D)** sienna4 depict genes with network membership |>| 0.6.

### Gene expression provides many unique and some shared routes to convergent behavioral evolution

To identify candidate genes that are robustly (i.e. independently across multiple analyses) associated with nest strategy and/or aggression, we integrated results across differentially expressed genes, individual gene- and network level PGLMMs, and concordant genes from at least 5 out of 10 RRHO comparisons (Table S11). For nest strategy-associated candidates, 27 out of 44 genes (61%) in the tan4 network recapitulated convergent individual genes in the PGLMM analysis, a strong signal that this gene set is consistently associated with the convergent evolution of obligate cavity-nesting. Out of the 221 differentially expressed genes shared among species pairs, 13 genes overlapped with nest-strategy associated individual PGLMM analyses and the tan4 network. We also identified 6 aggression-associated genes that overlapped with nest-strategy associated analyses, including 3 with concordant RRHO genes, 2 with individual PGLMM analyses, and 1 gene, TAF1B, which overlapped with both the tan4 network and individual PGLMM analyses. Altogether, these results suggest that repeated use of a small core set of genes may have contributed to the evolution of obligate cavity nesting and associated changes in behavior.

To place our results in a broader context, we compared our list of aggression-associated genes to prior studies (Data S12), including those on the brain’s response to an aggressive challenge (7, 54) and those that are known to mediate aggression in diverse species, including humans (22, 55). Using a permutation analysis, we found significant overlap between our 79 aggression-associated genes and another study’s network of 290 genes that were sensitive to experimental competition in the tree swallow brain (SI §11, Data S12, Table S12, Figure S19; (54)). Of the seven genes in common between these datasets (Table S13), *DPF3, PLXND1*, and *ZIC4* are involved in brain development, *CHN2* is associated with schizophrenia, *SPHKAP* is associated with neuroblastoma and apoptosis, and *GDF10* promotes neural repair after stroke (48), again linking aggression to key elements of brain function.

## Discussion

Striking phenotypic similarities among distantly related organisms exposed to similar ecological pressures yields critical insights into the mechanisms of evolution. However, investigations of the proximate mechanisms by which natural selection generates evolutionary change are often limited to traits with a simple and well-understood genetic basis (2, 3, 56), and less is known about the underpinnings of convergence in complex behavioral traits. A particularly unresolved question is the relative contributions of repeated vs. independent molecular changes to convergent behavioral evolution. Across five avian families with independent origins of obligate secondary cavity-nesting, we find behavioral convergence in territorial aggression, but limited transcriptomic convergence in the brain involving a small, core set of genes. These results highlight how complex behaviors can arise from largely independent evolution, punctuated by a few specific cases of repeated molecular changes in response to shared ecological pressures.

Across ten species, those with obligate cavity-nesting strategies exhibited greater physical aggression, though all species were present and responsive to the simulated intruder. Among females, aggression was particularly high for obligate cavity nesters, consistent with the hypothesis that female aggression is adaptive during competition for nesting territories (27, 37, 40, 57). Females were aggressive towards decoys of both sexes, whereas males were more aggressive towards their own sex. Conspecific aggression may serve multiple functions, most likely nest site defense or mate guarding (27, 58), considering the timing of stimulated intrusions during the early spring period of territorial establishment before egg laying. Our results complement a macroevolutionary study finding that cavity-nesters display more territorial behaviors against heterospecifics (59), underscoring multiple contexts linking territorial aggression with a cavity-nesting strategy.

Despite behavioral convergence, we did not observe higher levels of testosterone in obligate cavity-nesters. Testosterone-focused hypotheses have dominated aggression-related research for decades (18), yet we find that testosterone does not explain species-level differences in female or male territorial aggression, at least when birds are sampled at a constitutive state, without a prolonged social challenge (40). Recent phylogenetic analyses have found that breeding season length and mating system predict testosterone levels across species (60), but these life history traits are similar across our focal species. Interspecific variation in testosterone does not track interspecific variation in territorial aggression, for either sex. This finding aligns with recent perspectives that variation in testosterone is context-dependent (61, 62), and that testosterone is but one of many potential mechanisms regulating aggression (63, 64).

Using multiple phylogenetically informed approaches, we find that the convergent evolution of obligate cavity-nesting is associated with a small set of convergently expressed genes in the brain, alongside a larger set of lineage-specific genes shared only by some families or species. With these quantitative phylogenetic approaches, we find changes that occur more than is expected due to shared evolutionary history and random chance. These patterns could be driven by expression evolution in either the obligate-cavity or open nesters, but we are currently unable to resolve this directionality, given that we did not explicitly construct ancestral states for expression values. RRHO analyses revealed striking patterns of expression concordance between family comparisons, but only 0.1% of orthologs (11 genes) were associated with obligate cavity nesting across all comparisons. Single-gene and network PGLM analyses likewise revealed a small set of convergently evolving genes (0.4%; ∼40 genes). These proportions are similar to a recent phylogenetic study on muscle gene expression and courtship behavior in birds (14). However, they represent less convergence than most other studies of brain gene expression and behavioral evolution (9, 13, 65), which report that 4-6% of expressed orthologs are convergently evolving alongside behavior. Though our study differs by the number of taxa and/or degree of evolutionary divergence, another key difference is our explicit phylogenetic approach to identify convergently evolving genes, which statistically eliminates shared patterns of gene expression that stem from common ancestry and more directly tests for convergence. Thus, ours and other phylogenetic approaches may reduce the number of genes inferred as convergently evolving. Altogether, our results suggest that the convergent evolution of complex behavioral phenotypes can be underlain by mostly independent, lineage-specific changes in gene expression, with some convergent expression in a small, core set of genes shared across species.

This constellation of changes in gene expression may be associated with obligate cavity-nesting or aggression, and our PGLMMs disentangle these effects directly. Genes associated with nest strategy or aggression exhibited uncoupled regulatory evolution, with differing degrees of convergence and low overlap in gene identity. These patterns suggest that there are diverse neuromolecular correlates of aggression, including many that differ among species. Notably, we also found significant gene overlap with our aggression-associated genes and a previously published socially-sensitive gene network (54), indicating concordance between short-term functional responses to competition and long-term evolutionary change in competitive environments. Though our study focused on genes constitutively expressed during territorial establishment, future work should examine convergence in genes whose expression is activated or suppressed by acute social challenges, particularly since such environmentally sensitive genes may also play a key role in behavioral evolution (66).

By including both females and males, our study also assesses key elements of sex-specific evolution. Across species, we find sex-biased gene expression, for which males had higher expression that females. In birds, males carry two copies of the Z chromosome, and dosage compensation of the Z chromosome may be incomplete or absent (67), leading to higher expression of Z-linked genes in males than females. This mechanism likely explains our results for the ‘red’ network of co-regulated genes, which had higher expression in males than in females. Higher expression in males could also stem from stronger sexual selection (68), or greater constraint on the evolution of reduced expression (i.e. floor effects). Our sex-specific analyses further evaluate the potential for sex-specific behavioral and mechanistic convergence. Consistent with this hypothesis, we found that female obligate cavity nesters are more aggressive than females of both facultative and open-cup nesting species – a pattern that was not recapitulated in males. We also identified two female-specific gene networks that correlated with aggression. These sex-specific patterns suggest that female aggression has evolved differently from male aggression in obligate cavity nesting species, again underscoring multiple regulatory routes to behavioral convergence.

In summary, convergent evolution provides a natural experiment to identify the genomic underpinnings of complex phenotypes, and our work highlights the power of comparative approaches to uncover these mechanistic bases of convergence (13, 14, 21). Though future experiments are needed to functionally validate the nesting- and aggression-associated patterns we identify, our study is among the first to use phylogenetically explicit models of brain gene expression to understand the evolution of a complex and continuously varying behavioral trait.

Our findings support the hypothesis that there are largely independent evolutionary routes to building an aggressive bird, layered atop a small set of convergently evolving genes.

## Methods

This work was approved by Indiana University Bloomington IACUC #18-004 and #21-003, and relevant federal (MB59069B) and state permits (IL: W20.6355, KY: SC1911001, IN: 18-030, 19-272, 2545). Unless otherwise noted, analyses used R version 4 (R-Core-Team 2019).

### Assay of aggression

For cavity-nesting species, we placed the conspecific decoy on the nestbox and hung a Bluetooth speaker nearby. For non-cavity-nesters, we located nest sites or observed individuals for at least an hour to determine where they spent their time. For facultative cavity nesting species, including Carolina wrens and house sparrows, some aggression assays were conducted on individuals for which the nest could not be located, and therefore the nest type could not be confirmed. We compared levels of physical aggression and distance from decoy between individuals with a known nestbox vs. these other individuals and found no significant differences between groups for either species (p > 0.19). Additional details on audio stimuli and decoys can be found in SI §2. We played a conspecific vocal lure to capture the attention of the focal individual and waited 30 s before beginning the 5-min aggression assay. We measured a suite of aggressive behaviors (see SI §2) and focused on the proportion of the trial spent physically attacking the decoy. We calculated a maximum attack score of 60, based on the number of 5-second (sec) intervals that contained any physical contact. To visualize this behavior, we converted attack scores to a proportion of the trial spent attacking (# intervals including attack/total # intervals x 100). We also measured distance from the focal individual to the decoy to confirm that all focal territory holders were present and engaged with the simulated intruder. We evaluated the effects of nest strategy, sex, decoy sex, and their interaction on physical attacks and distance from the decoy using phylogenetic linear mixed models (details below and in SI §9). For downstream analyses, we include attack averages among species and sex categories as a fixed effect, referred to simply as ‘aggression.’

### Sample collection

For brain gene expression, we euthanized individuals with an anaesthetic overdose of isoflurane and rapid decapitation. We collected trunk blood into heparinized BD Microtainers (#365965). Using tools cleaned with RNAse-away and 95% ethanol, we immediately dissected out whole brains and flash froze on powdered dry ice within 8 min 26 sec ± 12 sec of euthanasia followed by storage at -80°C. Some additional hormonal samples for were obtained from barn swallows via brachial venipuncture (n = 4 females, n = 8 males), and sex was confirmed from blood via PCR (69). Blood was stored on ice packs in the field and later centrifuged for 10 min at 10,000 rpm. Plasma was separated and stored at -20°C. See SI §3 for details on enzyme immunoassays.

Immediate collections occurred an average of 18 min and 25 sec after the assay started. There was no relationship between latency to sampling and testosterone (Pearson’s correlation: r = 0.031, p = 0.84; Figure S3B), or brain gene expression (r |<| 0.32, p > 0.13). For the facultative cavity-nesters, we also found no difference in testosterone between individuals nesting in nestboxes compared with individuals whose nest type was unknown, for either species (p > 0.51), and no difference in brain gene expression for the tan4 eigengenes (p > 0.88).

### Brain dissection and RNA isolation

For 6 to 7 individuals per sex per species, we dissected whole brains into functional regions following (70). We removed the cerebellum, hindbrain, optic chiasm, optic tecta, and hypothalamus to the depth of the anterior commissure. Gene expression analyses focused on the ventromedial telencephalon (VmT), which contains nodes in the vertebrate social behavior network (47) including the extended medial amygdala, bed nucleus of the stria terminalis, and lateral septum (Figure S5). We collected this region by removing ∼1mm of the ventromedial portion of the caudal telencephalon. We extracted total RNA using Trizol (Invitrogen), and resuspended RNA in UltraPure water. Tape Station Bioanalyzer (Agilent) on 121 samples showed RIN = 8.56 ± 0.05 (mean ± SE).

### Sequencing, alignment, and mapping

We submitted total RNA to the Indiana University Center for Genomics and Bioinformatics, using paired end sequencing (Illumina NextSeq500, 75 cycle sequencing module) to generate an average of ∼28.6 million reads per sample (Data S1). We used Trinity version 2.13.2 (71) to assemble transcriptomes per species, which we aligned to the zebra finch (*Taeniopygia guttata*) proteome (NCBI assembly GCF_003957565.2_bTaeGut1.4.pri_protein.faa) (72) using tblastn v. 2.2.9. See SI §4 for additional bioinformatics details on alignment, mapping, and downstream analyses. We identified 10,672 orthologous genes with high confidence in all 10 species.. We normalized counts (Data S2) and performed differential gene expression using DESeq2 (73) (SI §5).

### Rank-Rank Hypergeometric Overlap

To identify shared transcriptomic patterns in obligate cavity-nesting species across pairs of families, we used the R package Rank-Rank Hypergeometric Overlap (RRHO) (74) (details in SI §8). RRHO uses normalized counts to rank genes based on the direction and magnitude of expression difference between groups, in our case each obligate cavity-nester and its species pair within the same family.

Hypergeometric tests evaluate overlap among these ranked lists, visualized as heatmaps of p-values from 100 permutations. The pipeline also identifies the set of genes with the highest degree of concordance for each comparison.

### Weighted Gene Co-Expression Network Analysis

We constructed gene networks for both sexes combined using WGCNA (75). We used the normalized counts from DESeq2 and filtered out genes with <15 norm counts in 90% of the samples. We generated a signed hybrid network by selecting a soft threshold power (β) = 6, in accordance with scale-free topology. We calculated a minimum network size of 30 and used a biweight midcorrelation (bicor) function. We merged modules in Dynamic Tree Cut using a threshold of 0.25. Genes with an absolute network membership value >0.6, which indicates high network connectivity, were assessed for enrichment of biological processes in PantherGO (76), using our list of 10,672 orthologs as a reference set. Networks of interest were visualized in Cytoscape v3.10.1 (77). See SI §8 for gene network analyses run separately on females and males.

### Phylogenetic Generalized Linear Mixed Models

Phylogenetic generalized linear mixed models (PGLMM) used MCMCglmm (78), with the phylogenetic covariance matrix from the consensus tree as a random effect (SI §9). Our consensus tree (Figure 1A) made use of the BirdTree.org (79) resource to obtain 1000 ultrametric trees from Ericson All Species. We constructed a majority rule consensus topology from this tree set using the *ape* (80) function ‘consensus’, and inferred branch lengths using the *phytools* (81) function ‘consensus.edges’ with the least squares method. For each PGLMM, we evaluated MCMC convergence using Heidelberger and Welch’s convergence diagnostic as implemented by the heidel.diag function in the R package coda.

To evaluate whether nest strategy predicted aggression, average distance from the decoy, and testosterone, we fit models with nest type, sex, and their interaction as fixed effects. Our aggression dataset contained many individuals that never attacked, and so we used a zero-inflated binomial distribution, with random effect and residual variance priors fixed at 1, the default burn-in period of 3000, and 2,000,000 iterations. For this aggression analysis, technical limitations in using this specific model with a zero-inflated binomial distribution restricted our ability to specify obligate cavity-nesting as the baseline factor level; therefore, we combined facultative nesting and open nesting into a single category to contrast with obligate cavity-nesting. For all other PGLMMs, we fit models containing either two (obligate vs. non-obligate) or three (obligate vs. facultative vs. open) factor levels (SI §9). For these other models, we fit a standard Gaussian model with no priors, the default burn-in, and 100,000 iterations.

To predict nest-strategy and aggression associated genes from our set of ∼10k orthologs, we included species/sex-level aggression sex, nest strategy, nest strategy by sex interaction and nest strategy by aggression interaction as fixed effects. After fitting a model for the log-expression of each gene, we applied a Benjamini-Hochberg false discovery rate correction in the *statsmodels* package (82) in Python version 3.7.11. Since our PGLMMs could not estimate p-values with precision lower than 1×10^−4^ (due to computational constraints), we set all raw p-values reported as less than this value to exactly 1×10^−4^. Since the Benjamini-Hochberg procedure relies on the relative ranks of raw p-values, rather than their magnitudes, this should not significantly affect our FDR correction.

To identify gene networks associated with nest strategy, sex, and their interaction we ran a similar phylogenetic analysis using the first eigengene value drawn from a weighted gene co-expression network analysis (SI §10).

Our PGLMM-related scripts are available at https://github.com/slipshut/CavityNesting/tree/main/PGLMM

### Functional interpretation

We inferred gene function based on genecards.org (48) and GO analyses. We inferred GO terms from *Homo sapiens* because its ontologies are orthologous to, but more complete than avian references.

We conducted an overrepresentation analysis of non-redundant biological process GO terms in PantherGO (76), using our list of orthologs as a reference set.

We also conducted a custom permutation analysis to evaluate the degree of overlap between our aggression-related genes and published genomic studies of aggression (7, 22, 54, 55) (Data S12; SI §11). Our enrichment scripts are available at https://github.com/slipshut/CavityNesting/tree/main/Enrichment

## Acknowledgements

For facilitating fieldwork, we thank the Indiana Department of Natural Resources, IU’s Research and Teaching Preserve, Thomas Rothfus and Chris Hagie at Therkilsden Field station, Auriel Fournier and Steve Havera at Forbes Field Station, Jeff Hoover, Nick Antonson, Hannah Scharf, and Angela Funk. For vocal stimuli, we thank Matt Wilkins, Dustin Reichard, Cara Krieg, and Elsa Chen. Thanks to Emmi Mueller for help with 3D printing; Sumitha Nallu and Jie Huang for TapeStation and library preparation; Emma Dossey Curole and Jace Kuske for assistance in the lab; Becca Young for assistance with data analysis; Amanda Clark for assistance with data visualization; and our reviewers for their feedback.

## Data Accessibility

Raw sequence reads and count data can be obtained from the Gene Expression Omnibus database (GEO accession number GSE244480). Supplementary information, including R scripts and trait data can be obtained from Github: https://github.com/slipshut/CavityNesting.

