## Supplementary Information for "Repeated behavioral evolution is associated with targeted convergence of gene expression in cavity-nesting songbirds"

This PDF file includes:

**Supporting Text**

SI § 1: Study species, study sites, and timing

SI § 2: Aggression assay

SI § 3: Testosterone enzyme immunoassay

SI § 4: Sequencing, alignment, and mapping

SI § 5: Differential gene expression within each avian family

SI § 6: Gene ontology enrichment analysis

SI § 7: Rank-rank hypergeometric overlap

SI § 8: Weighted gene co-expression networks

SI § 9: Phylogenetic generalized linear mixed models

SI § 10: WGCNA PGLMM – three level analysis of nest strategy

SI § 11: Permutation analysis: comparing aggression-associated genes to other candidates

**Tables S1-S14**

Table S1: Model coefficient estimates from PGLMM for physical attacks

Table S2: Model coefficient estimates from PGLMM for distance from decoy

Table S3: Model coefficient estimates from PGLMM for testosterone

Table S4: Differentially expressed genes between species pairs, for each family

Table S5: RRHO concordance permutation test for each pairwise family comparison

Table S6: Genes shared across all 10 family comparisons in RRHO

Table S7: PGLMM on nest strategy and aggression associated genes

Table S8: Global PGLMM on WGCNA networks that included both sexes

Table S9: Female-only PGLMM on WGCNA networks

Table S10: Male-only PGLMM on WGCNA networks

Table S11: Shared genes across analyses

Table S12: Overlap analysis of candidate gene datasets with aggression-associated genes

Table S13: Shared genes between aggression-related genes and Bentz brown network

**Figures S1-S17**

Figure S1: Aggression relative to the sex of the decoy, by sex and nest strategy

Figure S2: Distance from the decoy, by sex and nest strategy

Figure S3: Testosterone levels across collection approaches

Figure S4: Testosterone levels by sex and species

Figure S5: Landmark regions of the ventromedial telencephalon brain dissection

Figure S6: Gene expression PCA from 10,672 orthologs, by species and sex

Figure S7: UpSet plot of differentially expressed genes between species pairs

Figure S8: Number of differentially expressed genes vs. divergence time for 5 species pairs

Figure S9: Randomly permuted dataset run in RRHO for both sexes

Figure S10: Female-only RRHO

Figure S11: Male-only RRHO

Figure S12. Number of shared genes for each family comparison in RRHO

Figure S13: WGCNA networks associated with sex, nest strategy, and aggression

Figure S14: For females, WGCNA networks associated with nest strategy and aggression

Figure S15: For males, WGCNA networks associated with nest strategy and aggression

Figure S16: Overlap between aggression-genes and candidate datasets, 2-level analysis

Figure S17: Overlap between aggression-genes and candidate datasets, 3-level analysis

Figure S18: Overlap between aggression gene sets from Bentz et al. 2021, 2-level analysis

Figure S19: Overlap between aggression gene sets from Bentz et al. 2021, 3-level analysis

**SI References**

**Additional supporting materials for this manuscript include the following (online):**

Data S1: Read mapping

Data S2: Gene expression normalized counts

Data S3: Differential gene expression for species pair

Data S4: RRHO most up and down-regulated genes

Data S5: Genes shared across at least 5 out of 10 families, using RRHO comparisons

Data S6: Genes associated with sex, nest strategy, and aggression, 2-level analysis

Data S7: Genes associated with sex, nest strategy, and aggression, 3-level analysis

Data S8: Convergent genes associated with nest strategy and aggression

Data S9: WGCNA gene info and eigengenes

Data S10: PGLMM WGCNA – 2-level analysis

Data S11: PGLMM WGCNA – 3-level analysis

Data S12: Candidate aggression genes from literature, for overlap analysis

#### 1. Study species, study sites, and timing

We focused on 10 North American songbird species (oscine Passeriformes) with well-characterized ecologies and life histories (Figure 1A). In the Swallows (Hirundinidae), this included tree swallows (*Tachycineta bicolor*), which are obligate secondary cavity-nesters (1) and barn swallows (*Hirundo rustica*), which build open-cup mud nests in human-made structures (2). In the Woodwarblers (Parulidae), this included prothonotary warblers (*Protonotaria citrea*), which are obligate secondary cavity-nesters (3), and yellow warblers (*Setophaga petechia*), which build open-cup nests in bushes (4). In the true Sparrows (Passeridae), this included two introduced species, Eurasian tree sparrows (*Passer montanus*), which are obligate secondary cavity-nesters (5), and house sparrows (*Passer domesticus*), which are facultative cavity-nesters that may nest in enclosed domed nests in dense foliage or human-made structures (6). In the Thrushes (Turdidae), this included Eastern bluebirds (*Sialia sialis*), which are obligate secondary cavity-nesters (7), and American robins (*Turdus migratorius*), which build open-cup nests in trees and shrubs (8). In the true Wrens (Troglodytidae), this included house wrens (*Troglodytes aedon*), which are obligate secondary cavity-nesters (9), and Carolina wrens (*Thryothorus ludovicianus*), which are facultative cavity-nesters that may build a cup or dome nest within loosely enclosed natural and artificial locations (10).

We sampled barn swallows, yellow warblers, house sparrows, Eastern bluebirds, American robins, house wrens, and Carolina wrens at multiple sites around Bloomington, IN, USA (39142 N, 86.602 W). We sampled tree swallows near Lexington, KY (38.104 N, 84.489 W), prothonotary warblers near Belknap, IL (37.313 N, 88.984 W), and Eurasian tree sparrows near Havana, IL (40.353 N, 90.022 W) and Armington, IL (40.349 N, 89.349 W), USA. We studied additional barn swallows near Monticello, IL (40.028 N, 88.573 W) and additional house sparrows near Havana, IL. Sampling dates ranged from mid-March to mid-May from 2018 – 2021, depending on the species and localities, and included COVID-19 impacted study seasons. We collected and processed data from species pairs within the same family during the same years.

To ensure all species were in the territorial establishment and breeding stage, we monitored eBird records (11) and observed our field sites directly and daily as pairs formed and began defending territories, evidenced by singing and/or displacing intruders. For both facultative and obligate cavity-nesting species, we also noted their nestbox contents because focal species claim their territory with a piece of nest material placed in the artificial cavity. Similarly, barn swallows reuse mud nests year after year, so we monitored nests for occupancy and signs of re-building with fresh mud. For other species, we observed pairs for multiple hours to determine territorial boundaries. Thus, some criteria used to predict breeding stage varied in a species-specific manner, but our observations suggest that all individuals were within the territorial establishment phase, which overlaps with the period of nest-building but occurs before any eggs were formed and laid. Indeed, when we confirmed breeding status by post-mortem inspection of the gonad(s), we noted that females had small white follicles and males had enlarged, white testes, consistent with this pre-breeding stage. No females had large yolky follicles that would have indicated fertility.

#### 2. Aggression assay

To assay territorial aggression, we simulated intrusion for 5-min using randomized combinations of 3-4 male or female taxidermic decoys and 3-5 conspecific vocalizations. We recorded behaviors from both the female and male in each territorial pair. Behavioral observations were conducted ~15m from the decoy using binoculars, and behaviors were spoken into an audio recorder and then transcribed manually. We waited ~10 min after setup to begin the audio stimulus, or sooner if the focal individual began attacking.

To ensure that the focal individuals noticed the decoy, we used an audio playback of a conspecific lure and then waited 30s before starting the trial. We measured behaviors including attacks, which involved physical contact with the decoy using the beak or feet, distance to decoy at 1m intervals, as well as flyovers, hovers, and songs. We primarily focused on attacks as overt expressions of aggression during territorial competition. We calculated a maximum attack score of 60, based on the number of 5-second intervals that contained any physical contact during the 5-minute assay. We converted these scores to a proportion of the trial spent attacking (# intervals including attack/ total # intervals x 100). For instance, a focal individual that attacked during the entire trial would have a maximum score of 60 intervals, and an attack score of 100. Hereafter, we refer to this attack score as ‘aggression.’ We also analyzed distance from the decoy to confirm that all focal territory holders were present and engaged with the simulated intruder, whether or not they were aggressively attacking. For additional behaviors that reflect less escalated aggressive displays, we counted each instance of a newly initiated flyover, hover, or song during the 5-min trial; however, we did not include these behaviors in downstream analyses.

Audio stimuli were played from a Bluetooth speaker (JBL Charge 3) that was visually obscured inside a black or beige stocking, depending on habitat or nestbox characteristics. Audio stimuli were sourced from Xeno Canto, the Macaulay library, and colleagues (12) (E. George, D. Reichard, K. Krieg, and E. Chen; with permission). Stimuli consisted of songs or calls, depending on species-specific territorial contexts. This included songs from both sexes for barn swallows (12, 13), Eastern bluebirds (14), and house wrens (15), and defensive calls from both sexes for tree swallows (16). For the remaining species, we were not aware of evidence that females sing, so we used songs from males and calls from females. Although there is evidence of song in female prothonotary warblers in other populations (17), we were unable to obtain recordings and so used female chip calls. None of the playback stimuli were recorded from our focal populations.

Decoys used in our behavioral assays were made from birds collected during our research, as well as carcasses salvaged in good condition, in accordance with our permits. Briefly, we prepared these decoys following (18) by separating the skin/feathers from the body, and then affixed it to a 3D printed model posed in a natural position. We glued on black beads for eyes. We standardized decoy and speaker placement across trials and species to a frequently accessed area within the bird’s territory, though this varied to some degree with species-specific factors including vegetation and nest location. For obligate and facultative cavity-nesting species, we placed the decoy on top of the nest box (for prothonotary warblers, Eurasian tree sparrows, house sparrows, Eastern bluebirds, house wrens, and Carolina wrens), or in front of the nestbox hole (for tree swallows). For open cup nesting species (yellow warblers and American robins), we observed the focal pair for at least one hour to determine where they spent most of their time, and then placed the decoy and speaker in this area, in a tree or bush ~0.5 m from the ground, or ~0.5m below an elevated perch. For barn swallows, we placed the decoy within 0.5 m of the mud nest, on a tripod or ladder. We did not test decoys from individuals collected at the same locality to avoid potential familiarity with the simulated intruder.

#### 3. Testosterone enzyme immunoassay

We extracted steroids from plasma samples using diethyl ether (3x extractions) and reconstituted in 250μL assay buffer. We measured free testosterone using a High Sensitivity Testosterone Enzyme Immuno- Assay kit (Enzo #ADI-900-176, Farmingdale, NY, USA) following (19). We used 50μL plasma from females, and 10μL from males. We calculated T concentration by comparing sample absorbance with the absorbance of the assay’s standard curve, with all samples run in duplicate (Gen5 curve-fitting software, Biotek EPOCH plate reader, Winooski, VT, USA). We ran all samples in duplicate. We re-ran samples that showed greater than 80% maximum binding at 20μL plasma to obtain values in the most sensitive part of the curve. We ran samples that showed less than 20% maximum binding with 10μL reconstituted in 500μL assay buffer. Each plate also contained 3 duplicates from a pool of previously extracted plasma, used to calculate variability within and across plates. Intra-plate CV was 5.53% ± 0.81 and inter-plate CV was 25.7%. This elevated inter-plate CV likely relates to the use of 4 different kit lots across 4 years of study. However, we balanced families, species, and sexes within a plate each year.

#### 4. Sequencing, alignment, and mapping

We submitted total RNA to the Indiana University Center for Genomics and Bioinformatics in two rounds – once in 2019, for species pairs in the Thrush and Swallow families, and once in 2021 for species pairs in the Sparrow, Wren, and Warbler families. Because we sequenced species pairs together, we did not anticipate batch effects that would impact downstream gene expression analyses. Library preparation was conducted with a TruSeq Stranded mRNA HT sample preparation kit (Illumina). Paired end sequencing was conducted using an Illumina NextSeq500 with a 75 cycle sequencing module. We generated an average of ~28.6 million reads for the barcoded RNA samples (Data S1).

We demultiplexed sequencing reads with bcl2fastq (v2.20.0.422). Reads from the 2019 sequencing were trimmed for adapters and low-quality bases using Trimmomatic v0.36 (20). Reads from the 2021 sequencing were trimmed using fastp v0.20.1 (21) with parameters “-l 17 --detect_adapter_for_pe -g -p”. For each species, we used Trinity v2.13.2 (22) to assemble transcriptomes from species-specific reads. We then aligned the assembled transcripts to the zebra finch proteome (NCBI assembly GCF_003957565.2_bTaeGut1.4.pri_protein.faa) (23). The species we sampled range in divergence time from zebra finch between 22-26 mya, so we have no reason to expect biased mapping (25).

We had a multi-step process to assign zebra finch proteins to each species’ transcripts, particularly for identifying candidate aggression-related genes (Data S12). We initially used legacy tblastn v2.2.9 (24) with parameters “-F "m L" -U T -e 1e-3 -a 4 -v 5” to map our species transcriptomes against all zebra finch proteins. We then used cd-hit to cluster proteins and chose a ‘primary’ protein. Straightforward cases involved assigning proteins to transcript hits with a 1:1 best match. For ambiguous cases, including ties and multiple matches, we assigned proteins using a custom script:

h[ttps://github.iu.edu/abuechle/GSF2285_Rosvall_CavityNesting/blob/master/transcriptome_assemblies/blastBaseIdentity.protFixAmbig.pl](https://urldefense.com/v3/__https://github.iu.edu/abuechle/GSF2285_Rosvall_CavityNesting/blob/master/transcriptome_assemblies/blastBaseIdentity.protFixAmbig.pl__;!!OToaGQ!pB7WSy4hCvqt4e0eCpxlGtbgJ97P6FJO-1Ibc7UkCLASbKD4Ca3t9eRau8z9SC6xzUTKINb5O3gnyHHzz0g$). This script contained a cascade of rules that assigned proteins based on varying aspects of the hit, including highest percent ID (75% or greater), as well as different degrees of query length, subject length, etc. Per species GTF files were then created to define regions of transcripts that match to zebra finch proteins.

We quantified the transcript abundance in each species by mapping reads to their respective transcriptome assemblies using Bowtie2 v(26). Reads were sorted and indexed using Samtools (27). Reads mapped to transcriptomes and assigned to relevant Zebra finch proteins were quantified using featureCounts from the Subread package v2.0.2z with the parameters “-O -M -p --countReadPairs -B --primary --fracOverlap .75 --largestOverlap (28). Approximately 5.83 million reads per sample were mapped to the zebra finch proteome, which account for ~20% of reads in each sample (Data S1). After alignment and mapping, we focused on counts for 10,672 orthologous genes identified with high confidence in all 10 species.

We ran a principal component analysis and found that individuals clustered by family, species, and sex for the PCA of overall gene expression (Figure S6). PCA revealed clustering by sex, species, and family (Figure S6). PC1 explained 19% of the variance in gene expression, and PC2 explained 14%. One sample, a female house sparrow, had a divergent expression pattern from other samples, and upon further investigation also had lower overall expression relative to other individuals in same species and sex, and so was excluded.

#### 5. Differential gene expression within each avian family

We performed differential gene expression analysis using the *DESeq2* package v1.32.0 in R/Bioconductor (R v4.1.1) (29). We fit data to a negative binomial generalized linear model with a fixed effect for nest strategy for each transcript and filtered based on a per-transcript Wald test statistic to identify valid significantly differentially expressed transcripts. For each of the 5 families, we compared means of gene expression between species pairs using log2foldchange. *P*-values were corrected for multiple-testing (via the Benjamini-Hochberg procedure; adjusted *p* ≤0.05). We determined relative rank abundance using the normalized read counts generated by *DESeq2* (Data S2). Differential expression results are available in Data S3 for each gene, and summaries for both sexes together and separately are available in Table S4. Differential expression of orthologous genes ranged from 31-43% for each species pair within a family, for each sex (Table S4). An UpSet plot revealed unique and shared patterns of differential gene expression between species pairs. Notably, there were 221 differentially expressed genes shared across all five families (Data S3, Figure S7), several hundred genes that showed differential expression in just one family, and thousands of genes that showed differential expression in two to four families. Differential gene expression generally increased with the degree of divergence between species pairs within each family (t = 2.57, p = 0.08, R^2^ = 0.83; Figure S8; Table S4).

Scripts are available at <https://github.com/slipshut/CavityNesting/tree/main/DEGs>

#### 6. Gene ontology enrichment analysis

We conducted five types of GO analyses: 1) enrichment of differentially expressed genes within species pairs, 2) enrichment of genes significantly predicted by nest strategy, sex, and their interaction, 3) enrichment of WGCNA module eigengenes significantly predicted by nest strategy, sex, and their interaction, 4) enrichment of RRHO shared genes between species pairs, for each of the 20 pairwise family comparisons, and 5) enrichment of genes shared between PGLMM and RRHO analyses.

#### 7. Rank-rank hypergeometric overlap

Rank-rank hypergeometric overlap (RRHO) uses normalized read counts (Data S2) generated from DESeq2; see SI §6. We assigned a signed rank to genes within a family comparison based on the equation -log10(p-value)* sign(log2foldchange). Genes with lower expression in the cavity-nesting species are assigned a negative rank, while those with higher expression are given a positive rank. Based on rank, we placed genes from pairs of family comparisons into bins. Each bin represents a set of 104 ranks that share the same direction and similar magnitude of expression differentiation between obligate cavity-nesters and their relatives with more flexible nest strategies. Within a rank bin, a hypergeometric test is used to evaluate significant overlap of the genes present from the paired family comparisons. A significant p-value indicates more shared genes than would be expected by chance between the two comparisons. We generated hypergeometric p-value heatmaps across all bins for each comparison. To evaluate the significance of the distribution of p-values in a heatmap, we used a permutation test with 100 permutations per comparison as implemented in the RRHO package (30).

Pooling both sexes together can mask sex-specific patterns of gene expression, so we ran RRHO separately for females and males. We generated a list of the most upregulated and most downregulated genes from each pairwise family comparison for females, males, and both sexes combined (Figure S12), by applying a significance cutoff to the concordance results in Figures 3, S10, and S11. From this list, we counted the number of genes that overlap across family comparisons, ranging from 2 to 10 (Figure 3a).

As with the pooled sex dataset, gene expression differences remained highly concordant when RRHO was conducted on each sex individually. Patterns in the RRHO results with both sexes combined generally reflect the pattern seen in the sex with the stronger signature of overlap for that comparison.

There were also differences between the sex-specific datasets in the direction of shared changes (Figure S10, Figure S11). In the female-specific RRHO analysis, we saw a higher degree of concordance among 2-4 families (Figure 3A). Female obligate cavity nesters generally had lower expression than non-obligate species (Figure S10), whereas male obligate cavity nesters generally had higher expression than species with more flexible nesting strategies, though this pattern was not consistent for every comparison (Figure S11). In females, shared patterns of concordance were especially strong for the comparisons of families that included an obligate vs. facultative cavity nester – the Sparrow and Wren comparisons had stronger overlap in their heatmaps (e.g. higher p-values) (Figure S10). There were no significantly enriched GO terms. Similarly, this family comparison had a longer list of both the most upregulated and most downregulated genes (Figure S11). In the male-specific RRHO results, shared patterns of concordance were also strongest between Sparrows and Wrens (Figure S11). There were no significantly enriched GO terms.

For the female-specific RRHO, there were 9 shared downregulated genes and 3 shared upregulated genes across all family comparisons (Table S6). For the male-specific RRHO, there was one shared downregulated gene and 10 shared upregulated genes across all family comparisons (Table S6). None of these shared genes were significantly enriched for any GO terms. Overall, there were very few genes shared across all pairs of comparisons, highlighting again that global concordance is low.

Scripts are available at <https://github.com/slipshut/CavityNesting/tree/main/RRHO>

#### 8. Weighted gene co-expression networks

In addition to both sexes combined, we also ran separate network construction for females and males separately (31). We used the normalized counts from DESeq2 and filtered out genes with <15 norm counts in 90% of the samples. We ran hierarchical clustering and a principal components analysis to identify outliers, and excluded one female house sparrow and one female Eurasian tree sparrow sample. We generated a signed hybrid network by selecting a soft threshold power - 8 for females, and 6 for males - in accordance with scale-free topology. We calculated a minimum module size of 30 and used a biweight midcorrelation (bicor) function. We merged modules in Dynamic Tree Cut using a threshold of 0.25. Genes with an absolute module membership value >0.6 and/or <0.6 were assessed for enrichment of biological processes in PantherGO (32), using our list of orthologs as a reference set. Networks of interest were visualized in Cytoscape v3.10.1 (33).

Scripts are available at <https://github.com/slipshut/CavityNesting/tree/main/PGLMM/WGCNA>

#### 9. Phylogenetic generalized linear mixed models

Accounting for phylogenetic relationships among species using phylogenetic generalized linear mixed models (PGLMM), we evaluated the effect of nest strategy on aggression, testosterone, and gene expression using either two levels (obligate vs. non-obligate) or three levels (obligate vs. facultative vs. open). The main text presents results from the two-level analysis, with non-obligate cavity nesters as the reference. Here, we describe the three-level model results.

Aggression and testosterone PGLMMs

For the three-level models, found significant effects of nest strategy, sex, decoy sex, and their interaction on aggression. Obligate cavity-nesters spent more time attacking the decoy, compared to species with open nesting strategies (PGLMM: p = 0.0021; Figure 1A; Table S1). Males attacked significantly more than females (PGLMM: p = 0.044), though sex also interacted with obligate cavity-nesting strategy (PGLMM: p = 0.0071), driven by higher aggression in obligate cavity-nesting females compared to other females (Figure 1B). We found a significant interaction between the sex of the territory holder and the sex of the decoy (PGLMM: p = 0.014), such that that female territory holders attack female and male decoys with similar levels of aggression, whereas male territory holders attack male decoys significantly more than female decoys (Figure S1; Table S1).

Distance from the decoy was not related to nest strategy (PGLMM: p > 0.30), sex (p = 0.10), nor their interaction (p > 0.34) (Table S2). Thus, all species were in visual and audio proximity to the simulated intruder.

Males had significantly higher levels of testosterone in circulation than females (PGLMM: p < 0.0001), but testosterone was not related to nest strategy nor the interaction between nest strategy and sex (p > 0.35; Figure 2, Table S3)

Altogether, these results from the 3-level models were qualitatively similar to the 2-level models.

Scripts are available at <https://github.com/slipshut/CavityNesting/tree/main/Behavior_Testosterone>

Gene expression PGLMMs

In our three-level PGLMM, there are two nest strategy comparisons with respect to expression: obligate vs. facultative and obligate vs. open. PGLMM analyses identified hundreds of genes whose expression was associated with our model terms (Table S7, Data S7). This included 278 genes associated with facultative cavity-nesting and 111 genes associated with open nesting (both relative to obligate cavity-nesting). Sex had the largest influence on expression variation, with 462 genes. Sex-associated genes also have higher expression in males, much more often than lower expression, consistent with the two-level model (Table S7). Furthermore, 92 genes were associated with aggression and 174 genes exhibited an interaction between nest strategy and aggression. 235 genes exhibited a sex-specific association with open or facultative nesting; the 136 genes associated with the interaction of sex with facultative nesting were significantly enriched for ATP metabolic process (GO:0046034) (Table S7). No other sets of genes had significant GO enrichment. In comparison to the two-level model, we found more genes associated with specific nest strategies. These patterns may result from more specific nest strategy assignment.

Like in the two-level models, we identified convergently evolving genes that differed in a consistent direction in at least three of five family comparisons (Data S8). 135 of 373 (36%) genes associated with nest strategy met this criterion. Several of these genes have connections to ATP and mitochondrial functions (*ATP1B1, ATP5ME, COX3, MRPL1, MRLP21,* and *MRPS34*), or known associations with behavioral phenotypes (*GRM5, TRMT1L, NT5C2, BCHE*). Fewer convergent genes interacted with nest strategy and aggression (18 of 169; 11%) or nest strategy and sex (23 of 229; 10%). These results reinforce the findings of our two-level models in the main text: genes associated with a main effect of nest strategy were more often convergent than genes associated with other variables.

We found some consistent overlap between genes in the 2-level and 3-level models. For example, convergent genes associated with the interaction of nest strategy and aggression include *UNC13B* and *SREK1* (Data S8). Convergent genes associated with the interaction of nest strategy and sex were also shared between the 2-level and 3-level approaches, including *PTPRO*, *LOC116806872*, *RELT*, and *SREK1.*

Scripts are available at <https://github.com/slipshut/CavityNesting/tree/main/PGLMM/expression>

Candidate genes related to sex steroid hormone signaling

In our study, the most salient predictor of testosterone was sex. Across species, on average, male birds had higher levels of testosterone than females. Similarly, genes encoding the synthesis and reception of testosterone differed by sex. Males had significantly higher expression of androgen receptor (*AR,* PGLMM: p = 0.01), aromatase (*CYP19A1,* PGLMM: p = 0.0062), 5α-reductase type 2 (*SRD5A2,* PGLMM = 0.00026), and 17beta-hydroxysteroid dehydrogenase type 4 (*HSD17B4,* PGLMM: p < 0.0001). Females had significantly higher expression of estrogen receptor 2 (*ESR2,* PGLMM: p = 0.00041). Of these genes, only HSD17B4 is on the Z chromosome. We did not find an association between these genes and obligate cavity nesting.

#### 10. WGCNA PGLMM – three level analysis of nest strategy

Here we describe results for the PGLMMs of the weighted gene co-expression networks, for three nest strategies – obligate cavity-nesting, facultative cavity-nesting, and open cup nesting. For the dataset with both sexes combined, 4 networks were significantly associated with nest strategy, aggression, sex, and their interactions: brown, darkgreen, red, and tan4 (Data S11, Table S8). As with the two-level models, the three-level model recovered the sex-associated red network and nest strategy-associated tan4 network (Figure S13A, Figure 3); additional trait-associated networks (brown and darkgreen) appear to be driven mainly by one putatively outlier species (Figure S13B,C). The brown network was significantly enriched for mitochondrial translation (Table S8). Other networks were not significantly enriched.

For the female-specific dataset, there were 3 networks associated with nest strategy and/or the interaction between nest strategy and aggression: royalblue, turquoise, and coral3 (Data S11, Table S9). These networks were again driven mainly by one putatively outlier species (Figure S14). Turquoise was significantly enriched for mitochondrial translation and oxidative phosphorylation (Table S9); other networks were not significantly enriched. There were also two gene networks associated with aggression in females: pink4 (Pearson’s correlation: r = 0.78, t = 9.34, df = 57, p < 0.0001) and sienna4 (r = -0.78, t = -9.54, df = 57, p < 0.0007) (Figure 4). Though neither of these networks were significantly enriched for any GO biological process terms, they did contain genes with known connections to aggression, for instance dopamine receptor DRD3 in pink4 and glutamate receptor GRIA2 in sienna4.

For the male-specific dataset, there were 4 networks associated with nest strategy and/or the interaction between nest strategy and aggression: blue, darkorange2, purple, and yellowgreen (Data S11, Table S10). Again, each of these trait-associated networks appear to be driven by one species (Figure S15). The blue network was significantly enriched for mtDNA translation and oxidative phosphorylation (Table S10).

Across each WGCNA, there were several networks significantly associated with the facultative nest strategy, but upon visual inspection they were driven by patterns in just one species: house sparrows or Carolina wrens. This included the brown and darkgreen networks for both sexes combined (Figure S13B,C Table S8), the royalblue, turquoise and coral3 networks for females (Figure S14A,B,C Table S9), and the blue, purple, and yellowgreen networks for males (Figure S15A,C,D Table S10). Of these, the brown, turquoise, and blue networks were enriched for biological processes relating to mitochondrial expression and function. In a study across mice, fish, and honeybees, the downregulation of oxidative phosphorylation genes was identified as a core process modulating neurogenomic responses to social challenge, indicating the importance of energy metabolism for territorial aggression (34).

Scripts are available at <https://github.com/slipshut/CavityNesting/tree/main/PGLMM/WGCNA>

#### 11. Permutation analysis: comparing aggression-associated genes to candidate genes

We conducted a custom permutation analysis to evaluate the degree of overlap between our aggression-genes and candidate genes that have been related to aggression in previous genomic analyses in human and rodent models (35), and captive fish (36), responses to social challenge in captive mice (34) and responses to experimental competition in wild birds (37) (Data S12). Since Bentz et al. focused on birds, we looked further at each set of differentially expressed genes and WGCNA network (MM > 0.6) associated with competition.

For the 2-level analysis, there were 79 aggression-related genes (Data S6). For the 3-level analysis, there were 92 aggression-related genes (Data S7). We first calculated the percentage of genes present in each of our candidate gene lists. We then randomly sampled the 79 (2-level) or 92 (3-level) genes without replacement from our set of 10,672 orthologs, and the percentage of genes from this resampled dataset that were represented in our candidate gene lists was also calculated. This procedure was repeated 10,000 times to generate a null distribution of enrichments for each candidate gene list. To test for significant enrichment of genes, we assigned rank *i* to the percent enrichment of the gene set against the list of 10,000 randomly sampled gene enrichments. A two-tailed p-value was then estimated using the following formula:

p = 1 – 2|0.5 - i|

This formula measures the deviation of our observed gene enrichment from the center of the resampled dataset, which has a rank of 5000. A significant p-value indicates that more or fewer genes from our list of genes are candidate behavioral genes than expected by chance. Custom R scripts are available at https://github.com/slipshut/CavityNesting/tree/main/Enrichment

For only the 3-level analysis, we found significant overlap between genes associated with aggression and a gene network associated with response to experimentally-induced territorial competition in another study of tree swallows (Table S12, Figure S19; Bentz et al. 2021). Of the seven genes in common between these datasets (Table S13), DPF3, PLXND1, and ZIC4 are involved in brain development, CHN2 is associated with schizophrenia, SPHKAP is associated with apoptosis and neuroblastoma, and GDF10 promotes neural repair after stroke.

### **SI Tables**

#### **Table S1**: Model coefficient estimates from Phylogenetic Linear Mixed Models (PGLMMs) for physical attacks, as measured by physical contact with a decoy during the 5-min aggression assay. Zi is the zero-inflated class. For the 2-level models, the baseline is obligate for main effect terms, and non-obligate for interaction terms. For the 3-level models, the baseline is open for main effect terms and facultative for interaction terms.

| **coefficients** | **post.mean** | **l -95% CI** | **u -95% CI** | **eff.samp** | **pMCMC** |
| --- | --- | --- | --- | --- | --- |
| **2-Level Nest Strategies** |  |  |  |  |  |
| Physical Attacks | -4.28 | -7.25 | -1.34 | 7353.51 | **0.01097** |
| Physical Attacks (zi) | -33.32 | -60.55 | -1.07 | 3.94 | **0.00655** |
| Nest Strategy (Non-Obligate) | -4.29 | -6.69 | -1.92 | 3809.52 | **0.00105** |
| Sex (Male) | 1.85 | -0.09 | 3.79 | 54657.15 | 0.05811 |
| Decoy Sex (Male) | -0.34 | -1.74 | 1.06 | 68149.38 | 0.63067 |
| Nest Strategy (Obligate) x Sex (Male) | -2.55 | -4.46 | -0.7 | 53667.08 | **0.00637** |
| Sex (Male) x Decoy Sex (Male) | 2.26 | 0.52 | 4.03 | 88591.04 | **0.01073** |
| **3-Level Nest Strategies** |  |  |  |  |  |
| Physical Attacks | -9.514 | -13.042 | -6.067 | 7642.746 | **0.00003** |
| Physical Attacks (zi) | -42.249 | -65.167 | -4.307 | 1.919 | **0.00072** |
| Nest Strategy (Facultative) | 2.163 | -1.945 | 6.198 | 2586.864 | 0.281 |
| Nest Strategy (Obligate) | 5.282 | 2.221 | 8.42 | 4504.635 | **0.00213** |
| Sex (Male) | 2.394 | 0.00845 | 4.766 | 62315.019 | **0.044** |
| Decoy Sex (Male) | -0.333 | -1.738 | 1.075 | 91004.59 | 0.642 |
| Nest Strategy (Obligate) x Sex (Male) | -3.024 | -5.331 | -0.799 | 74418.802 | **0.00711** |
| Nest Strategy (Open) x Sex (Male) | -1.121 | -4.29 | 2.026 | 59303.199 | 0.481 |
| Sex (Male) x Decoy Sex (Male) | 2.183 | 0.445 | 3.965 | 88715.232 | **0.01414** |

#### **Table S2:** Model coefficient estimates from Phylogenetic Linear Mixed Models (PGLMMs) for distance from decoy during the 5-min aggression assay. Models were run with 2 nest strategies (obligate vs. non-obligate cavity-nesting) and with 3 nest strategies (obligate cavity, facultative cavity, and open cup nesting). For the 2-level models, the baseline is non-obligate cavity. For the 3-level models, the baseline is obligate cavity.

| **Coefficients** | **post.mean** | **l-95% CI** | **u-95% CI** | **eff.samp** | **pMCMC** |
| --- | --- | --- | --- | --- | --- |
| **2-Level Nest Strategies** |  |  |  |  |  |
| Intercept | 5.4643 | 2.9214 | 7.9813 | 9700 | **0.00144** |
| Nest Strategy (Obligate) | -1.476 | -3.8743 | 0.9726 | 9700 | 0.22392 |
| Sex (Male) | 0.3534 | -1.4584 | 2.138 | 9700 | 0.70371 |
| Nest Strategy (Obligate) x Sex | 1.0811 | -1.4082 | 3.4928 | 9366 | 0.3901 |
| **3-Level Nest Strategies** |  |  |  |  |  |
| Intercept | 3.9925 | 1.2784 | 6.4632 | 9700 | 0.00701 |
| Nest Strategy (Facultative) | 1.1959 | -2.2711 | 4.3114 | 9700 | 0.4532 |
| Nest Strategy (Open) | 1.6429 | -1.4483 | 4.8598 | 9700 | 0.29814 |
| Sex (Male) | 1.4009 | -0.2672 | 3.1057 | 9700 | 0.10412 |
| Nest Strategy (Facultative) x Sex (Male) | -1.4911 | -4.5875 | 1.6196 | 9700 | 0.33918 |
| Nest Strategy (Open) x Sex (Male) | -0.6274 | -3.6754 | 2.5242 | 9700 | 0.69423 |

#### **Table S3:** Model coefficient estimates from PGLMM for testosterone. Models were run with 2 nest strategies (obligate vs. non-obligate cavity-nesting) and with 3 nest strategies (obligate cavity, facultative cavity, and open cup nesting). For the 2-level models, the baseline is non-obligate cavity. For the 3-level models, the baseline is obligate cavity.

| **Coefficients** | **post.mean** | **l-95% CI** | **u-95% CI** | **eff.samp** | **pMCMC** |
| --- | --- | --- | --- | --- | --- |
| **2-Level Nest Strategies** |  |  |  |  |  |
| Intercept | -0.68207 | -0.90066 | -0.46649 | 10645 | **<0.0001** |
| Nest Strategy (Obligate) | 0.02489 | -0.18095 | 0.21944 | 9700 | 0.799 |
| Sex (Male) | 1.11809 | 0.98293 | 1.25455 | 9700 | **<0.0001** |
| Nest Strategy (Obligate) x Sex (Male) | -0.02052 | -0.21309 | 0.16758 | 9700 | 0.846 |
| **3-Level Nest Strategies** |  |  |  |  |  |
| Intercept | -0.64969 | -0.85172 | -0.45655 | 9700 | **<0.0001** |
| Nest Strategy (Facultative) | 0.07451 | -0.19652 | 0.33557 | 9700 | 0.572 |
| Nest Strategy (Open) | -0.10393 | -0.3311 | 0.11606 | 9700 | 0.352 |
| Sex (Male) | 1.10045 | 0.96776 | 1.23072 | 9700 | **<0.0001** |
| Nest Strategy (Facultative) x Sex (Male) | 0.07608 | -0.18856 | 0.3479 | 9700 | 0.569 |
| Nest Strategy (Open) x Sex (Male) | -0.01497 | -0.22991 | 0.20409 | 9700 | 0.891 |

#### **Table S4:** Differentially expressed genes between species pairs, for each family, for both sexes together and separately, out of 10,672 orthologs, along with divergence time (mya) between species pair within each family.

| **Family** | **Divergence** | **# DEGs both sexes** | **# DEGs female** | **# DEGs male** |
| --- | --- | --- | --- | --- |
| Sparrows | 7.14 | 3551 | 3351 | 3337 |
| Woodwarblers | 9.14 | 3673 | 3357 | 3388 |
| Wrens | 15.5 | 3564 | 3577 | 3265 |
| Swallows | 19.5 | 4058 | 4083 | 3649 |
| Thrushes | 21.9 | 4545 | 4307 | 4440 |

#### **Table S5:** RRHO concordance permutation test for each pairwise family comparison. Comparisons with randomly permuted differential expression values were run to establish a null expectation, relative to comparisons with empirical differential expression values for both sexes combined, as well as females and males separately.

| **Comparison** | **Both Sexes**  **Randomized**  **p-value** | **Both Sexes Empirical**  **p-value** | **Females**  **Empirical**  **p-value** | **Males**  **Empirical**  **p-value** |
| --- | --- | --- | --- | --- |
| Sparrows vs. Swallows | 0.37 | < 0.01 | < 0.01 | < 0.01 |
| Sparrows vs. Thrushes | 0.81 | < 0.01 | < 0.01 | < 0.01 |
| Sparrows vs. Warblers | 0.25 | < 0.01 | < 0.01 | < 0.01 |
| Sparrows vs. Wrens | 0.44 | < 0.01 | < 0.01 | < 0.01 |
| Swallows vs. Thrushes | 0.51 | < 0.01 | < 0.01 | < 0.01 |
| Swallows vs. Warblers | 0.82 | < 0.01 | < 0.01 | < 0.01 |
| Swallows vs. Wrens | 0.04 | < 0.01 | < 0.01 | < 0.01 |
| Thrushes vs. Warblers | 0.97 | < 0.01 | < 0.01 | < 0.01 |
| Thrushes vs. Wrens | 0.83 | < 0.01 | < 0.01 | < 0.01 |
| Warblers vs. Wrens | 0.88 | < 0.01 | < 0.01 | < 0.01 |

#### **Table S6:** Genes shared across all 10 family comparisons (RRHO)

| **Comparison** | **Direction** | **Gene** |
| --- | --- | --- |
| Both sexes | Down | AFAP1L1 |
| Both sexes | Down | ATP6V0E1 |
| Both sexes | Down | B3GALT6 |
| Both sexes | Down | CTNND1 |
| Both sexes | Down | DDX56 |
| Both sexes | Down | EMC3 |
| Both sexes | Down | NPTX1 |
| Both sexes | Down | RGS19 |
| Both sexes | Down | WIPF2 |
| Both sexes | Up | LOC121470538 |
| Both sexes | Up | PQBP1 |
| Females | Down | ANKRD44 |
| Females | Down | CYTL1 |
| Females | Down | DDX56 |
| Females | Down | DLX5 |
| Females | Down | EFNB1 |
| Females | Down | HINT2 |
| Females | Down | LOC100217958 |
| Females | Down | SDC1 |
| Females | Down | WDR24 |
| Females | Up | LOC105758557 |
| Females | Up | LOC121470538 |
| Females | Up | METTL26 |
| Males | Down | ELOVL7 |
| Males | Up | PIK3R3 |
| Males | Up | LOC115494570 |
| Males | Up | LOC115494818 |
| Males | Up | LOC115495273 |
| Males | Up | LOC121470538 |
| Males | Up | METTL26 |
| Males | Up | PPP1R14D |
| Males | Up | RAC2 |
| Males | Up | RBM15B |
| Males | Up | SF3B5 |

#### **Table S7:** Phylogenetic Linear Mixed Models on individually expressed genes associated with nest strategy, aggression, and sex. Models run with 2 nest strategies (obligate vs. non-obligate cavity-nesting) had non-obligate as the baseline. Models run with 3 nest strategies (obligate cavity, facultative cavity, and open cup nesting) had obligate as the baseline. The total number of significantly associated genes for each model is listed, as well as genes that had higher or lower expression relative to the baseline. Gene Ontology terms listed were significantly associated with the total number of genes for each model.

| **Model Term** | **Total** | **Higher** | **Lower** | **Gene Ontology terms** |
| --- | --- | --- | --- | --- |
| **2-Level Nest Strategies** |  |  |  |  |
| Nest Strategy (Obligate) | 234 | 127 | 107 | None statistically significant |
| Nest Strategy (Obligate) x Aggression | 62 | 19 | 43 | None statistically significant |
| Nest Strategy (Obligate) x Sex (Male) | 76 | 33 | 43 | None statistically significant |
| Aggression | 79 | 47 | 32 | None statistically significant |
| Sex (Male) | 510 | 372 | 138 | cellular metabolic process (GO:0044237) |
| **3-Level Nest Strategies** |  |  |  |  |
| Nest Strategy (Facultative) | 278 | 85 | 193 | None statistically significant |
| Nest Strategy (Open) | 111 | 45 | 66 | None statistically significant |
| Nest Strategy (Facultative) x Aggression | 82 | 59 | 23 | None statistically significant |
| Nest strategy (Open) x Aggression | 92 | 39 | 53 | None statistically significant |
| Nest Strategy (Facultative) x Sex (Male) | 136 | 29 | 107 | ATP metabolic process (GO:0046034) |
| Nest Strategy (Open) x Sex (Male) | 99 | 53 | 46 | None statistically significant |
| Sex (Male) | 462 | 364 | 98 | None statistically significant |

#### **Table S8.** Global Phylogenetic Linear Mixed Models (PGLMMs) on Weighted Gene Coexpression Network Analyses (WGCNA) that were significantly associated with nest strategy and/or sex. Models run with 2 nest strategies (obligate vs. non-obligate cavity-nesting) had non-obligate as the baseline. Models run with 3 nest strategies (obligate cavity, facultative cavity, and open cup nesting) had obligate cavity-nesting as the baseline. Networks with significant fixed effects are shown here. Gene Ontology terms listed were significantly associated with genes that had a network membership |>| 0.6.

| **Network** | **PGLMM Significant Terms** | **Number of Genes** | **Number of Genes <**  **-0.6** | **Number of Genes > 0.6** | **High Network Membership Genes** | **Gene Ontology Terms** |
| --- | --- | --- | --- | --- | --- | --- |
| **2-Level Nest Strategies** | | | | | | |
| Red | Sex (Male) | 193 | 9 | 114 | 123 | None statistically significant |
| Tan4 | Nest strategy (Obligate) | 44 | 16 | 19 | 35 | None statistically significant |
| **3-Level Nest Strategies** | | | | | | |
| Brown | Nest strategy (Facultative), and all interactions with nest strategy (Facultative) | 303 | 21 | 203 | 224 | mtDNA translation (GO:0032543) |
| Dark Green | All interactions with nest strategy (Facultative) | 102 | 68 | 18 | 86 | None statistically significant |
| Red | Sex (Male) | 193 | 9 | 114 | 123 | None statistically significant |
| Tan4 | Nest strategy (Open) | 44 | 16 | 19 | 35 | None statistically significant |

#### **Table S9.** Female-only Phylogenetic Linear Mixed Models (PGLMMs) on Weighted Gene Coexpression Network Analyses that were significantly associated with aggression and/or nest strategy. Models run with 2 nest strategies (obligate vs. non-obligate cavity-nesting) had non-obligate as the baseline. Models run with 3 nest strategies (obligate cavity, facultative cavity, and open cup nesting) had obligate as the baseline. Networks with significant fixed effects are shown here. Gene Ontology terms listed were significantly associated with genes that had a network membership |>| 0.6.

| **Network** | **PGLMM Significant Terms** | **Number of Genes** | **Number of Genes < -0.6** | **Number of Genes > 0.6** | **High Network Membership Genes** | **Gene Ontology Terms** |
| --- | --- | --- | --- | --- | --- | --- |
| **2-Level Nest Strategies** | | | | | | |
| Pink4 | Aggression | 54 | 16 | 34 | 50 | No statistically significant results |
| Sienna4 | Aggression | 54 | 15 | 28 | 43 | No statistically significant results |
| Royalblue | Nest strategy (Obligate) x Aggression | 114 | 30 | 67 | 97 | No statistically significantly results |
| **3-Level Nest Strategies** | | | | | | |
| Royalblue | Nest strategy (Facultative) x Aggression | 114 | 30 | 67 | 97 | No statistically significantly results |
| Turquoise | Nest strategy (Facultative); Nest strategy (Facultative) x Aggression | 297 | 14 | 212 | 226 | mtDNA translation (GO:0032543), oxidative phosphorylation (GO:0006119) |
| Coral3 | Nest strategy (Facultative) | 55 | 11 | 39 | 50 | No statistically significantly results |

#### **Table S10.** Male-only Phylogenetic Linear Mixed Models (PGLMMs) on Weighted Gene Coexpression Network Analyses that were significantly associated with aggression and/or nest strategy. Models were run with 3 nest strategies (obligate cavity, facultative cavity, and open cup nesting), with obligate as the baseline. There were no models for 2 nest strategies with significant fixed effects. Gene Ontology terms listed were significantly associated with genes that had a network membership |>| 0.6.

| **Network** | **PGLMM Significant Terms** | **Number of Genes** | **Number of Genes < -0.6** | **Number of Genes > 0.6** | **High Network Membership Genes** | **Gene Ontology Terms** |
| --- | --- | --- | --- | --- | --- | --- |
| **3-Level Nest Strategies** | | | | | | |
| Blue | Nest strategy (Facultative), nest strategy (Facultative) x Aggression | 332 | 36 | 217 | 253 | mtDNA translation (GO:0032543), oxidative phosphorylation (GO:0006119) |
| Darkorange2 | Nest strategy (Facultative) x Aggression | 75 | 13 | 51 | 64 | No statistically significant results |
| Purple | Nest strategy (Facultative) | 151 | 28 | 97 | 125 | No statistically significant results |
| Yellowgreen | Nest strategy (Facultative) x Aggression | 84 | 17 | 59 | 76 | No statistically significant results |

#### **Table S11.** Shared genes among convergent nest strategy-related and aggression-associated individual genes, the tan4 WGCNA network, concordant RRHO genes, and differentially expressed genes shared across families.

| **Gene** | **Gene Description** | **Convergent Individual PGLMM** | **Tan4 Network PGLMM** | **Aggression PGLMM** | **RRHO 5 of 10** | **DEG Shared** |
| --- | --- | --- | --- | --- | --- | --- |
| AARS1 | alanine--tRNA ligase, cytoplasmic | X | X |  |  |  |
| ACTR3B | actin-related protein 3B isoform X2 | X | X |  |  |  |
| ADI1 | acireductone dioxygenase 1 | X |  |  |  | X |
| BCHE | cholinesterase isoform X1 | X | X |  |  | X |
| C14H7orf50 |  |  | X |  | X |  |
| CDADC1 | cytidine and dCMP deaminase domain-containing protein 1 | X | X |  |  | X |
| CLCC1 | chloride channel CLIC-like protein 1 isoform X3 | X | X |  |  | X |
| COPB1 | COPI coat complex subunit beta 1 |  | X |  |  | X |
| COX7C | cytochrome C oxidase subunit 7C |  |  | X | X |  |
| CTBP2 | C-terminal binding protein 2 |  |  | X | X |  |
| DLGAP2 | disks large-associated protein 2 isoform X17 | X | X |  |  |  |
| EML6 | echinoderm microtubule-associated protein-like 6 isoform X1 | X | X |  |  | X |
| GPCPD1 | glycerophosphocholine phosphodiesterase GPCPD1 isoform X5 | X | X |  |  |  |
| GPR180 | integral membrane protein GPR180 isoform X1 | X | X |  |  |  |
| GTDC1 | glycosyltransferase like domain containing 1 | X |  |  |  | X |
| HIGD1A | HIG1 hypoxia inducible domain family member 1A | X |  | X |  |  |
| HSPA2 | heat shock-related 70 kDa protein 2 | X | X |  |  |  |
| LOC100224313 | C-terminal-binding protein 2 | X |  |  |  | X |
| LOC115495400 | uncharacterized LOC115495400 | X | X |  |  |  |
| LRRC7 | leucine rich repeat containing 7 | X |  |  |  | X |
| MOGAT1 | 2-acylglycerol O-acyltransferase 1 | X | X |  |  | X |
| NIBAN1 | protein Niban 1 | X | X |  |  |  |
| NIT2 | omega-amidase NIT2 | X | X |  |  |  |
| NPTN | neuroplastin | X | X |  |  |  |
| NSUN4 | 5-methylcytosine rRNA methyltransferase NSUN4 | X | X |  |  |  |
| NT5C2 | cytosolic purine 5'-nucleotidase isoform X4 | X | X |  |  |  |
| POGLUT1 | protein O-glucosyltransferase 1 | X | X |  |  | X |
| POT1 | protection of telomeres protein 1 isoform X3 | X | X |  |  |  |
| PROM1 | prominin-1 isoform X6 | X | X |  |  |  |
| PTPDC1 | protein tyrosine phosphatase domain containing 1 | X |  |  |  | X |
| RNASEH2B | ribonuclease H2 subunit B | X |  | X |  |  |
| SLC25A24 | calcium-binding mitochondrial carrier protein SCaMC-1 | X | X |  |  |  |
| SLC45A1 | solute carrier family 45 member 1 |  | X |  |  | X |
| SLC7A6OS | solute carrier family 7 member 6 opposite strand | X |  |  |  | X |
| SLC8A3 | sodium/calcium exchanger 3 isoform X6 | X | X |  |  |  |
| TAF1B | TATA box-binding protein-associated factor RNA polymerase I subunit B isoform X1 | X | X | X |  |  |
| TBC1D30 | TBC1 domain family member 30 isoform X3 | X | X |  |  |  |
| THAP1 | THAP domain containing 1 |  |  | X | X |  |
| TXNRD3 | thioredoxin reductase 3 | X | X |  |  | X |
| XYLB | xylulose kinase isoform X1 | X | X |  |  |  |

#### **Table S12**: Overlap analysis of candidate behavioral gene datasets with aggression-associated genes. For each dataset, “observed overlap” is the proportion of observed aggression genes that overlap with each candidate gene dataset. “Mean randomized overlap” is the average proportion of overlapping genes from 1000 gene lists randomly sampled from our full list of ~10,000 orthologs. P-values are calculated by measuring where the observed overlap falls in the distribution of randomized overlap values (Figures S12-S15). A significant p-value means the observed overlap value is more extreme than 95% of randomized overlap values (in either direction).

|  | **2 nest strategies** | | | **3 nest strategies** | | |
| --- | --- | --- | --- | --- | --- | --- |
| **Candidate gene dataset** | **Observed overlap** | **Mean randomized overlap** | **p-value** | **Observed overlap** | **Mean randomized overlap** | **p-value** |
| Filby (36) | 0 | 0.00157 | 0.2342 | 0 | 0.00154 | 0.2682 |
| Rittschof (34) | 0 | 0.0127 | 0.729 | 0.0217 | 0.0126 | 0.2148 |
| Zhang & James (35) | 0.139 | 0.1018 | 0.2088 | 0.0869 | 0.1017 | 0.7922 |
| Pooled (non-bird) (38, 39, 40) | 0.139 | 0.113 | 0.345 | 0.1086 | 0.1129 | 0.9398 |
| Bentz pooled (37) | 0.038 | 0.0452 | 0.9748 | 0.0978 | 0.0459 | **0.018** |
| Bentz HYPO Day0 DEG | 0 | 0.0007 | 0.1102 | 0 | 0.0007 | 0.1298 |
| Bentz HYPO Day2 DEG | 0 | 0.0047 | 0.6302 | 0.0109 | 0.0046 | 0.1338 |
| Bentz VMT Day0 DEG | 0 | 0.0033 | 0.4636 | 0 | 0.0032 | 0.5126 |
| Bentz VMT Day2 DEG | 0 | 0.0014 | 0.204 | 0 | 0.0014 | 0.2444 |
| Bentz VMT WGCNA Blue | 0.0126 | 0.0133 | 0.5656 | 0.0109 | 0.0134 | 0.6914 |
| Bentz HYPO WGCNA Brown | 0.0253 | 0.0205 | 0.4412 | 0.0761 | 0.0209 | **0.0008** |
| Bentz HYPO WGCNA Green | 0 | 0.0044 | 0.5862 | 0 | 0.0043 | 0.6552 |

#### **Table S15.** Shared genes between aggression-related genes and Bentz et al. 2021 hypothalamus WGCNA brown module

| **Gene** | **Gene Description** |
| --- | --- |
| CHN2 | Chimerin 2, rho GTPase-activating protein 3 |
| DPF3 | Double PHD fingers 3 |
| GDF10 | Growth differentiation factor 10 |
| MYO3B | Myosin IIIB |
| PLXND1 | Plexin D1 |
| SPHKAP | SPHK1 interactor, AKAP domain containing |
| ZIC4 | Zinc finger protein of the cerebellum 4 |

### **SI Figures**

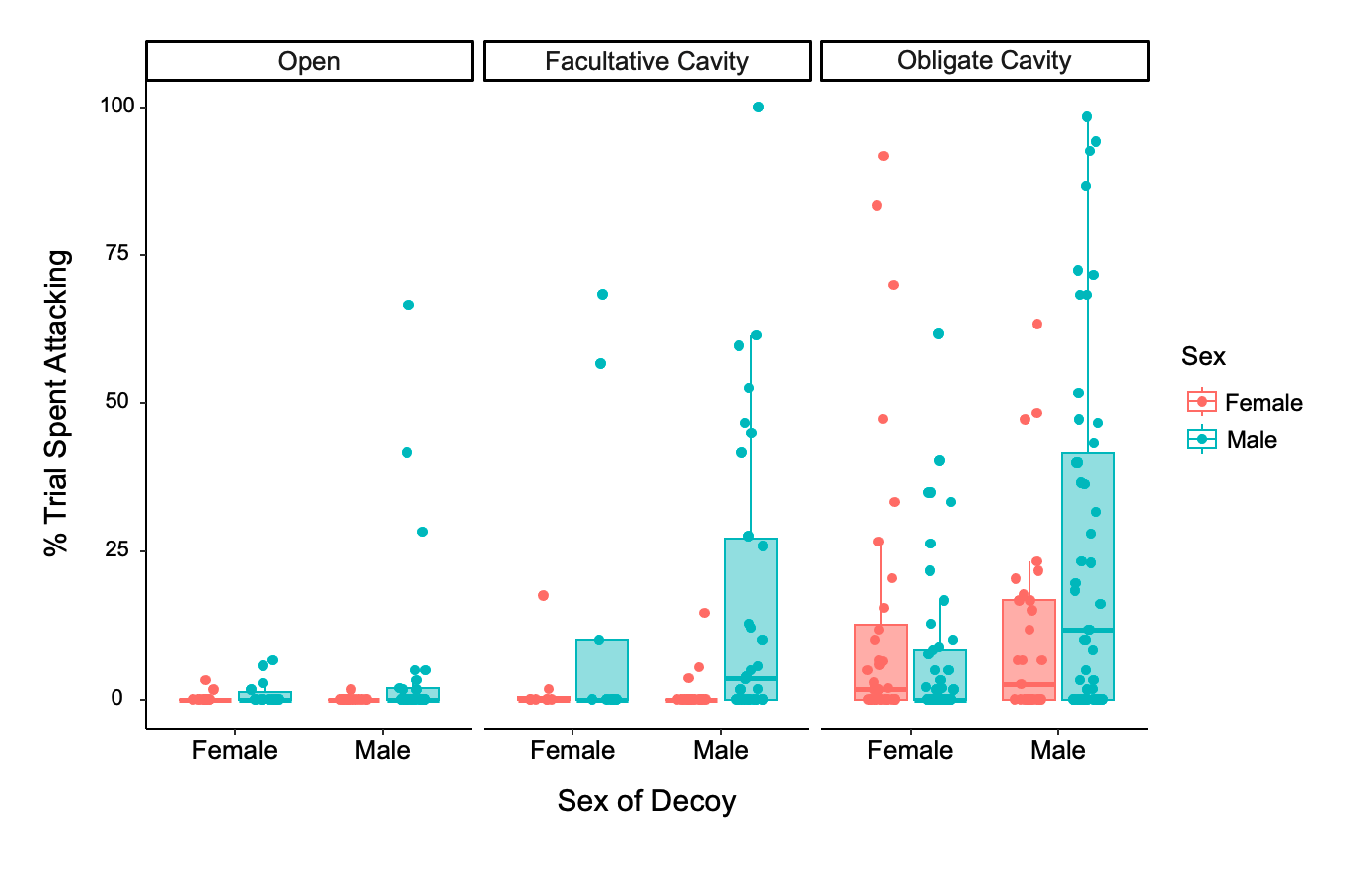

#### **Figure S1:** Aggression relative to the sex of the decoy for females and males from species with open, facultative cavity, and obligate cavity-nesting strategies. Box and whisker plots depict 10th, 25th, 50th, 75th, and 90th percentiles.

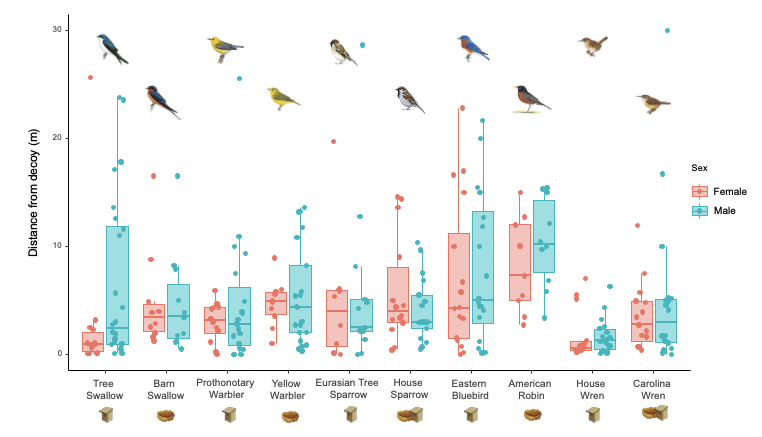

#### **Figure S2:** Distance from decoy for females and males from species with open, facultative cavity, and obligate cavity-nesting strategies. Box and whisker plots depict 10th, 25th, 50th, 75th, and 90th percentiles.

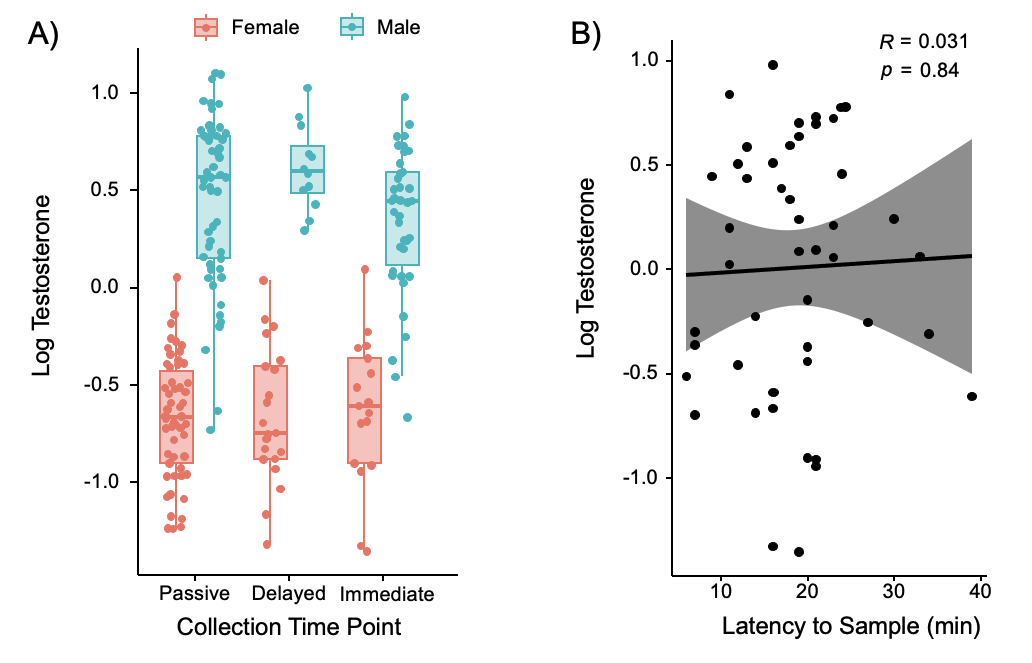

#### **Figure S3:** Testosterone levels across a) collection approaches and b) time to blood sampling from initiation of aggression assay. Box and whisker plots depict 10th, 25th, 50th, 75th, and 90th percentiles.

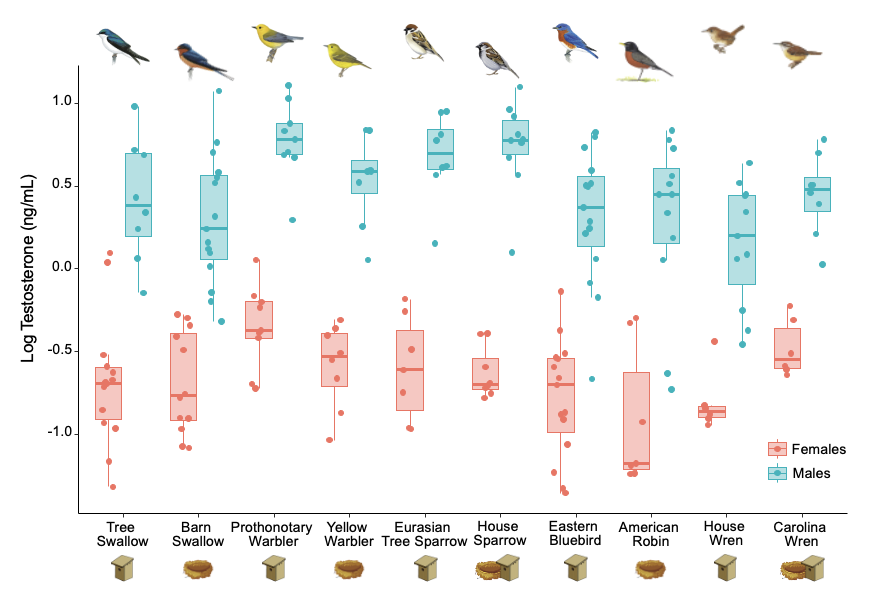
***Figure S4:*** *Testosterone levels from blood plasma across species and sexes. Illustrations reproduced with permission from Lynx Edicions. Box and whisker plots depict 10th, 25th, 50th, 75th, and 90th percentiles.*

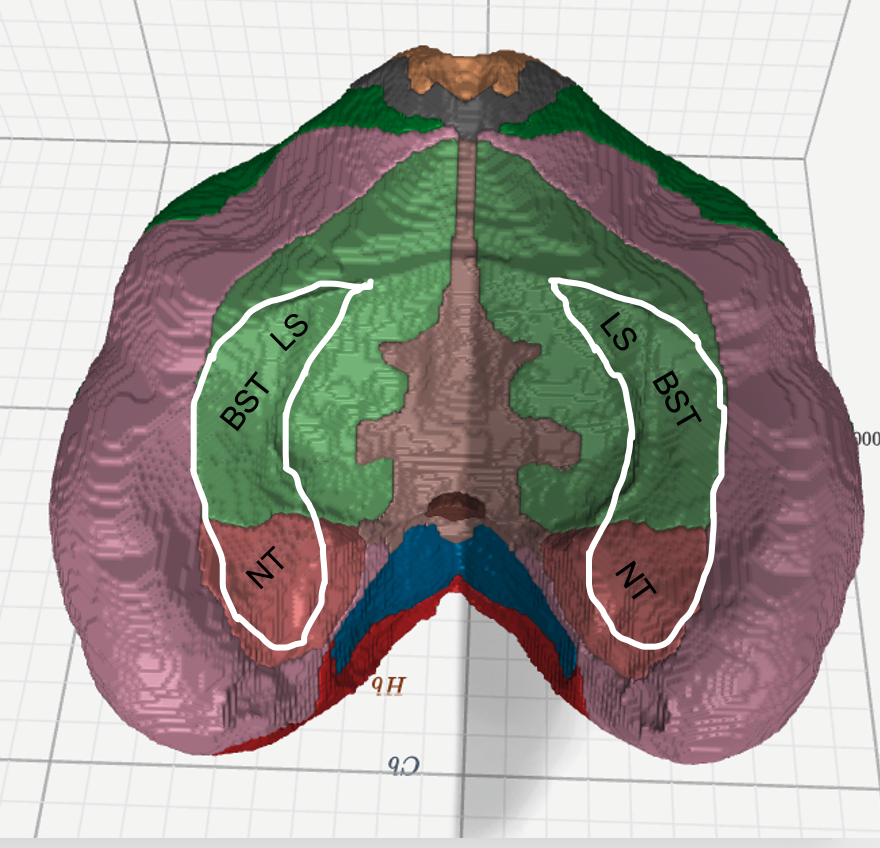

***Figure S5:*** *Landmark regions of the ventromedial telencephalon, including lateral septum (LS), bed nucleus of the stria terminalis (BST), and medial amygdala (NT). 3D canary atlas (38), image from https://timsainburg.com/pages/bird-brain-atlases.html*

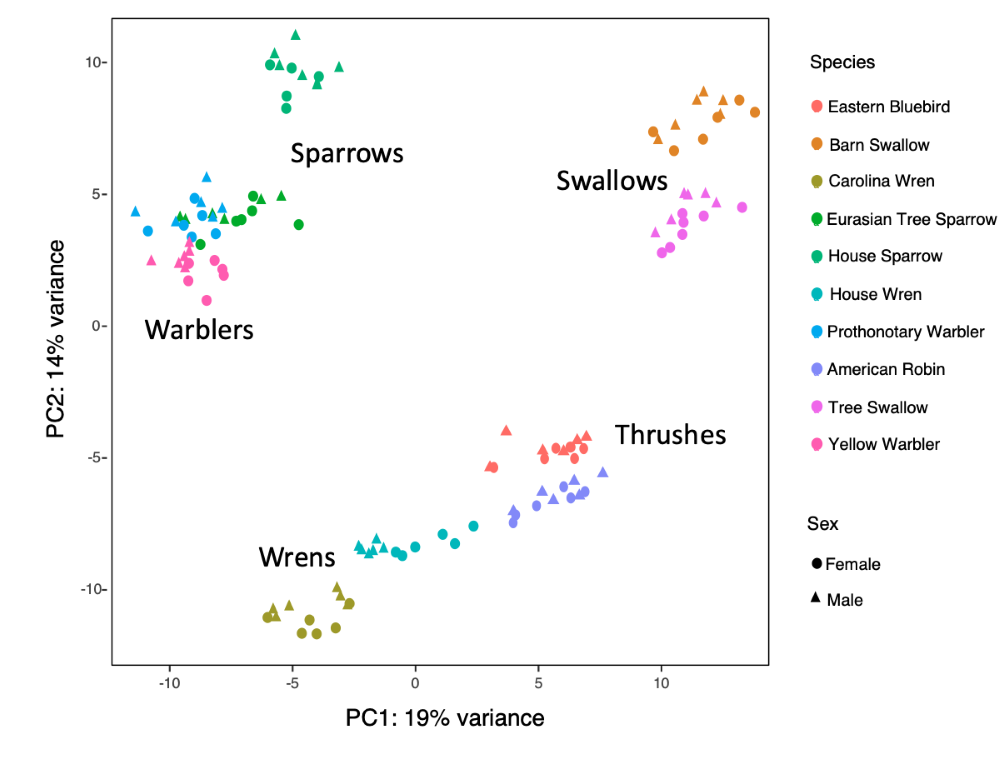

***Figure S6:*** *Gene expression PCA from 10,672 orthologs, by species and sex*

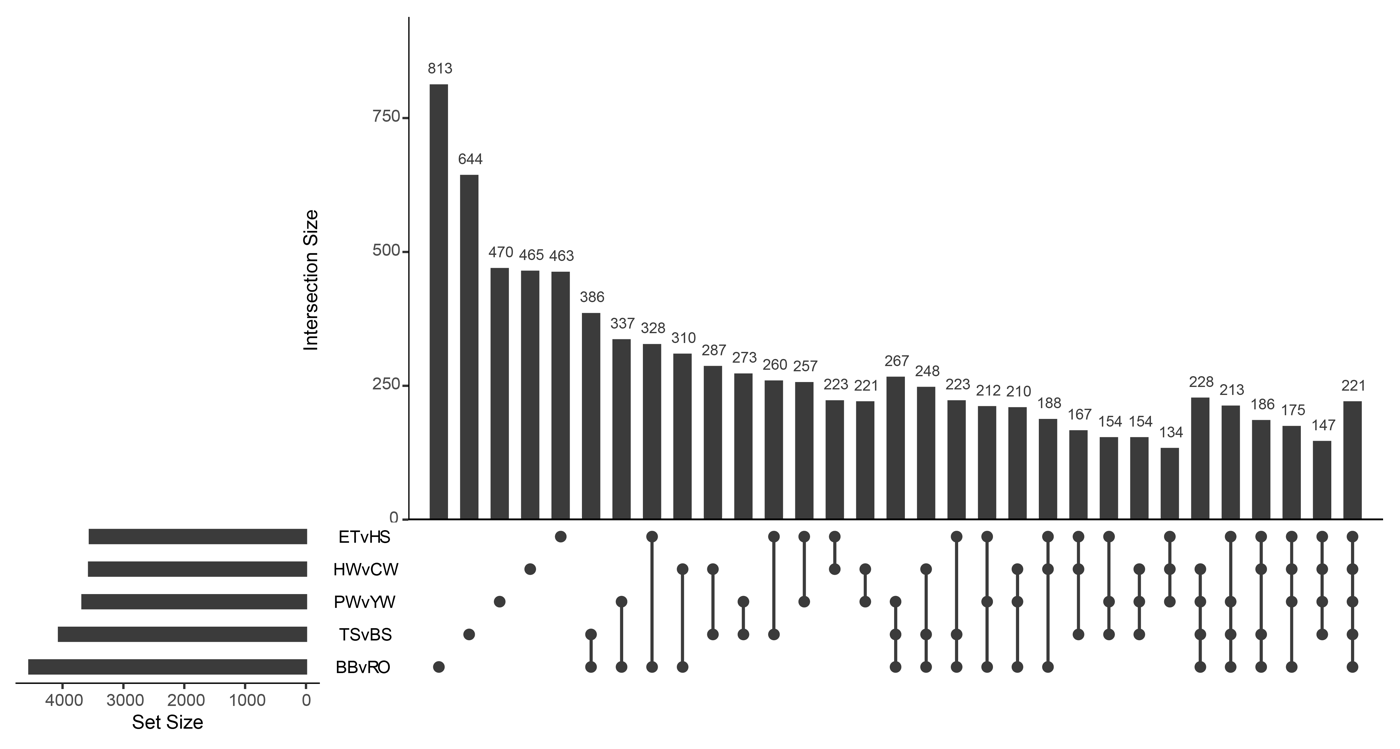

#### **Figure S7:** UpSet plot depicting the number of unique and shared differentially expressed genes between house wrens (HW) and Carolina wrens (CW), Eurasian tree sparrows (ET) and house sparrows (HS), prothonotary warblers (PW) and yellow warblers (YW), tree swallows (TS) and barn swallows (BS), and Eastern bluebirds (BB) and American robins (RO). Set size is the full list of differentially expressed genes, whereas the intersections size focuses on unique and shared differentially expressed genes with a log2foldchange > |0.5| and adjusted p-value < 0.05. Black dots on the x-axis represent whether these genes are present or absent in that set.

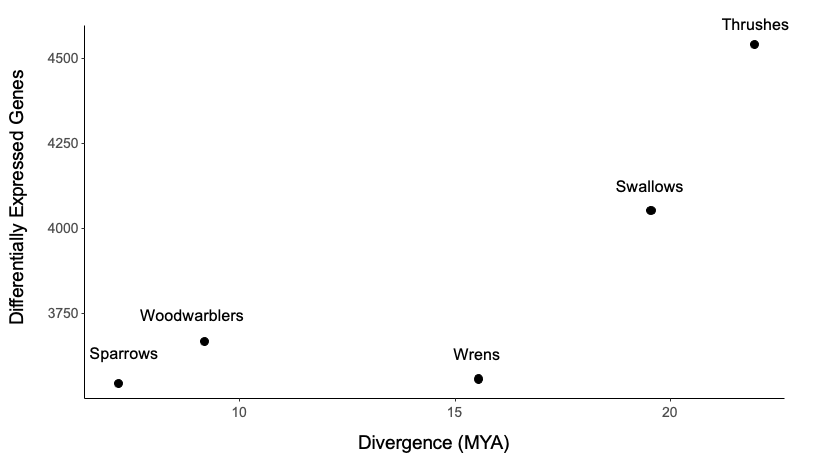

#### **Figure S8:** Divergence time in millions of years between species pairs vs. differentially expressed genes with a log2foldchange > |0.5| and adjusted p-value < 0.05.

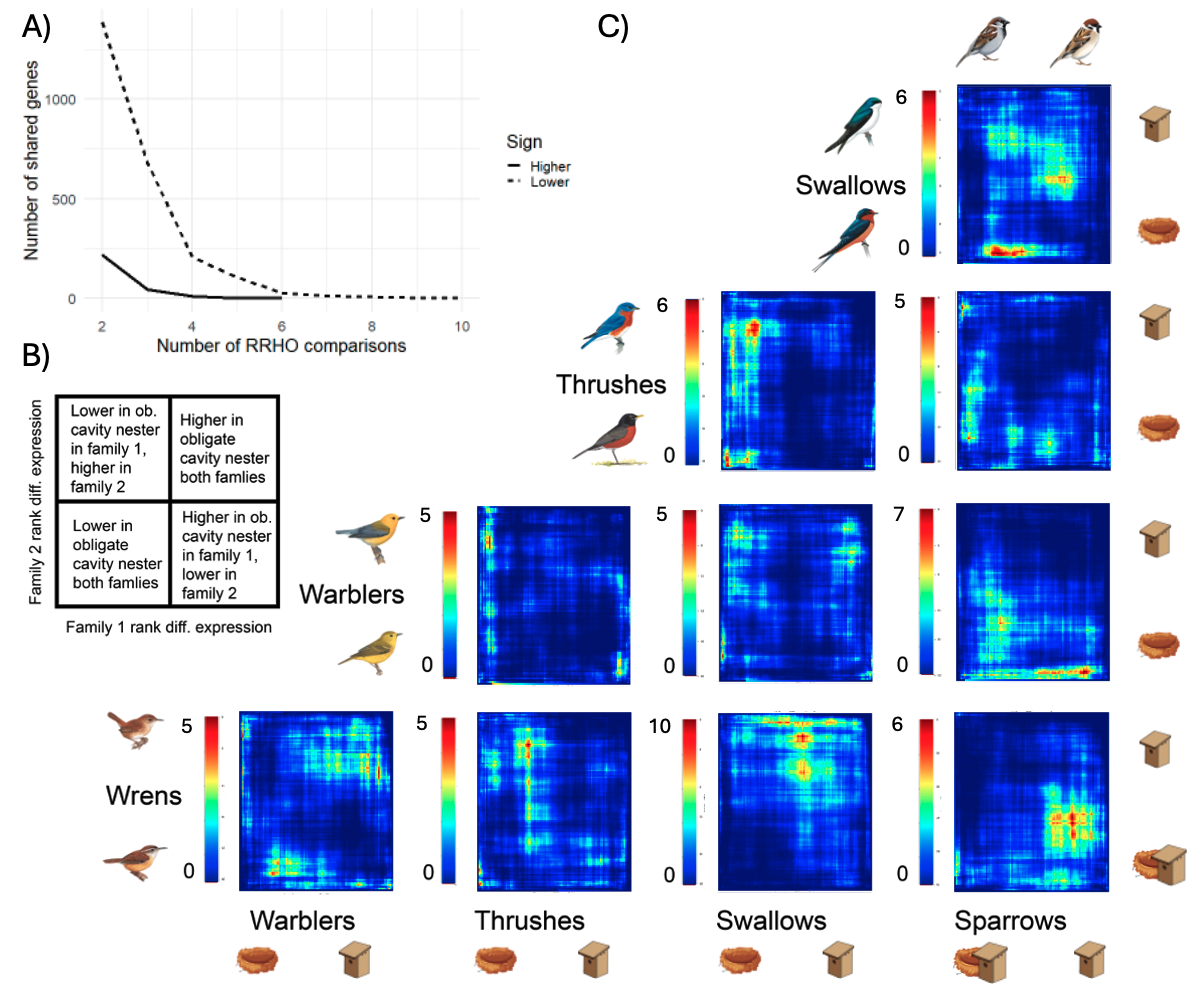

#### **Figure S9:** RRHO on randomly permuted data reveals a lack of concordance in differential expression among each family comparison. Heatmap colors reflect adjusted hypergeometric -log(p-value). Nestboxes indicate obligate cavity nesters, nests represent open nesters, and both together represent facultative cavity nesters. **(A)** Number of concordantly expressed genes shared across family comparisons (i.e. across heatmaps) for both sexes combined. Solid lines indicate higher expression in obligate cavity-nesters, dashed lines indicate lower expression **(B)** Key for interpreting individual heatmaps. Each quadrant of the key corresponds to a quadrant of an individual heatmap, shown in **(C)** Each pixel within the heatmaps contains two sets of approximately 100 genes being compared between the two families; the color legend indicates the log p-value of the overlap between these gene sets, with a higher value indicating stronger overlap.

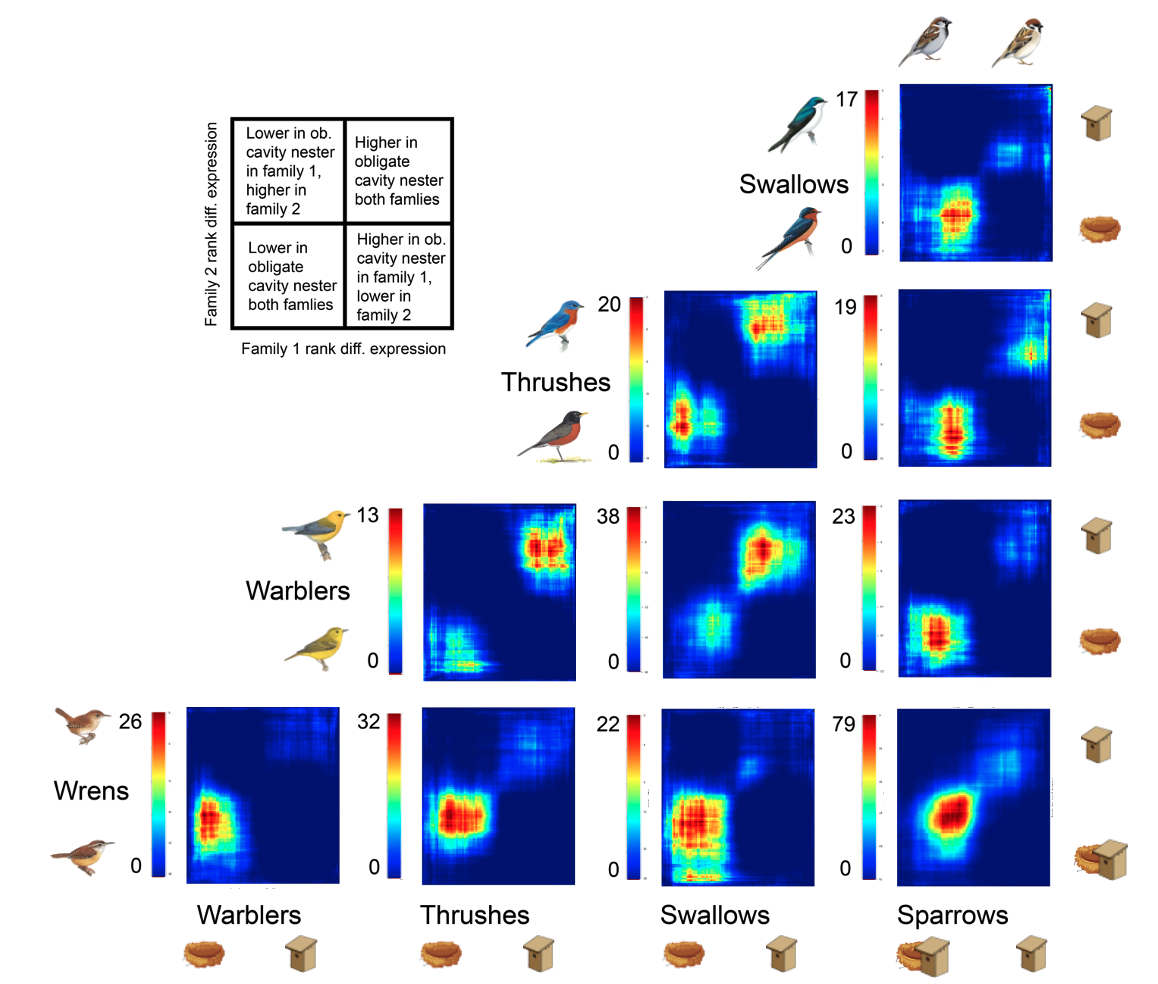

#### **Figure S10:** Female-only RRHO. Concordance in differential expression of all gene orthologues among each family comparison. Heatmap colors reflect adjusted hypergeometric -log(p-value). Nestboxes indicate obligate cavity nesters, nests represent open nesters, and both together represent facultative cavity nesters. Illustrations reproduced with permission from Lynx Edicions.

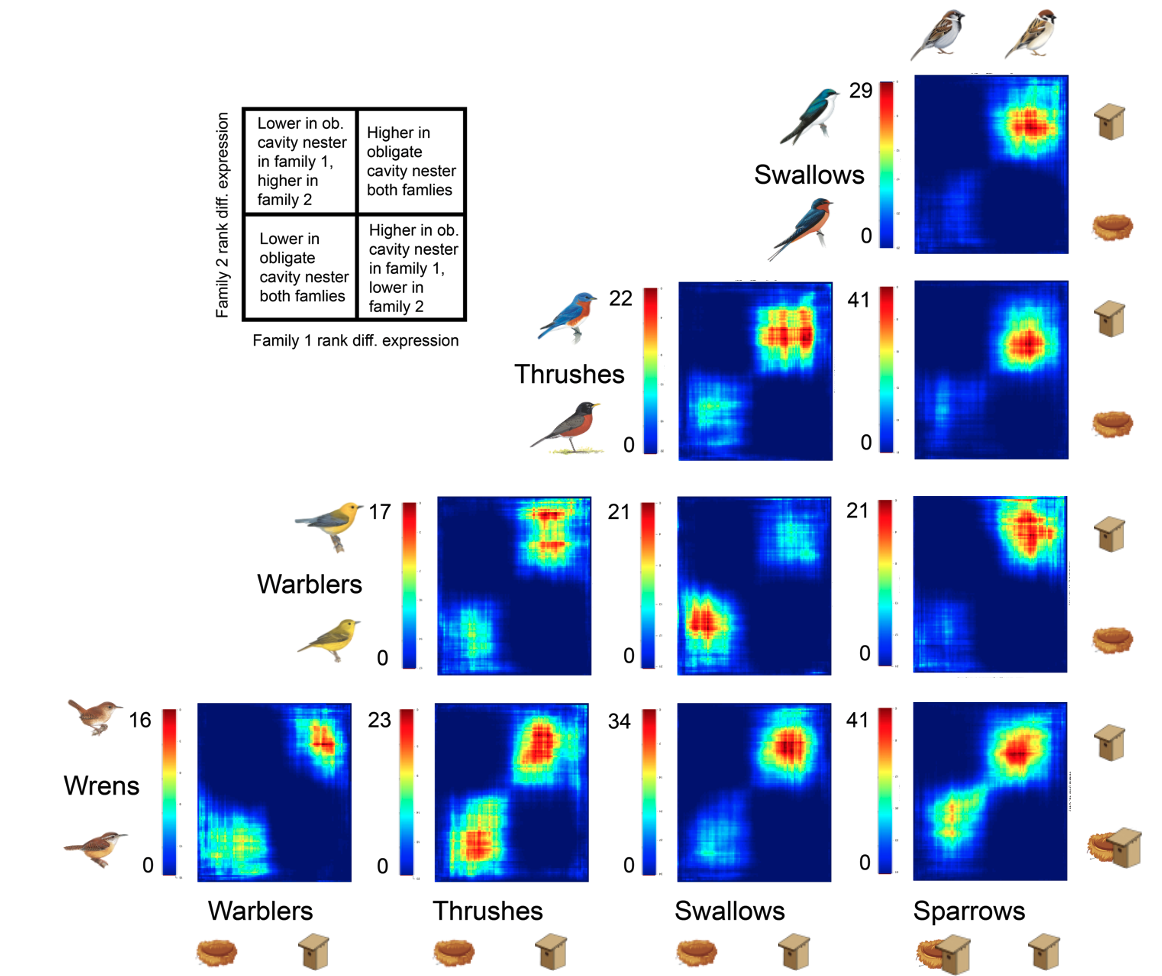

#### **Figure S11:** Male-only RRHO. Concordance in differential expression of all gene orthologues among each family comparison. Heatmap colors reflect adjusted hypergeometric -log(p-value). Nestboxes indicate obligate cavity nesters, nests represent open nesters, and both together represent facultative cavity nesters. Illustrations reproduced with permission from Lynx Edicions.

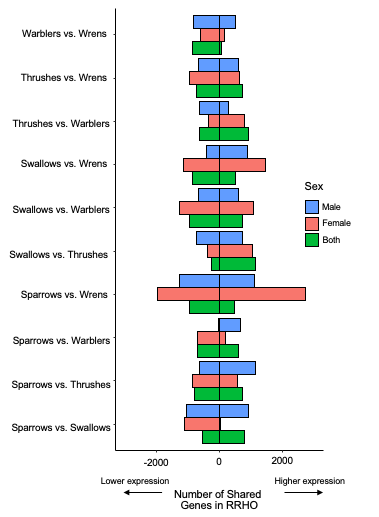

#### **Figure S12**. Number of shared genes that were most down-regulated or most up-regulated for each family comparison in RRHO. Individual gene info is available in Data S4.

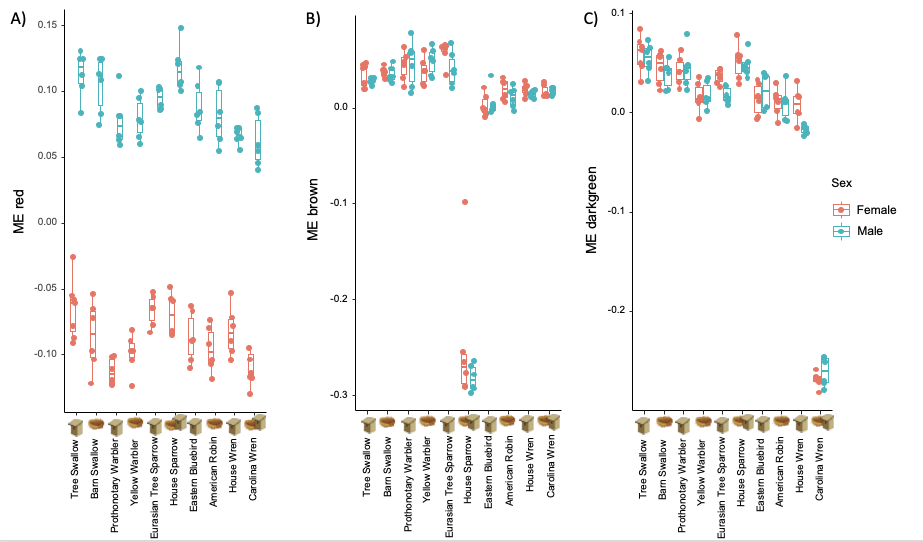

#### **Figure S13.** For both sexes, WGCNA module eigengenes associated with sex, nest strategy, and aggression. A) Gene expression in the red module was significantly associated with sex across species, for both the 2- and 3-strategy models. B) Gene expression in the brown module was significantly associated with facultative cavity-nesting and all interactions with the facultative cavity strategy (sex, aggression), for the 3-strategy model. This pattern appears driven by house sparrows. C) Gene expression in the darkgreen module was significantly associated with the interaction of facultative cavity-nesting with sex and aggression, for the 3-strategy model. This pattern appears driven by Carolina wrens. Box and whisker plots depict 10th, 25th, 50th, 75th, and 90th percentiles.

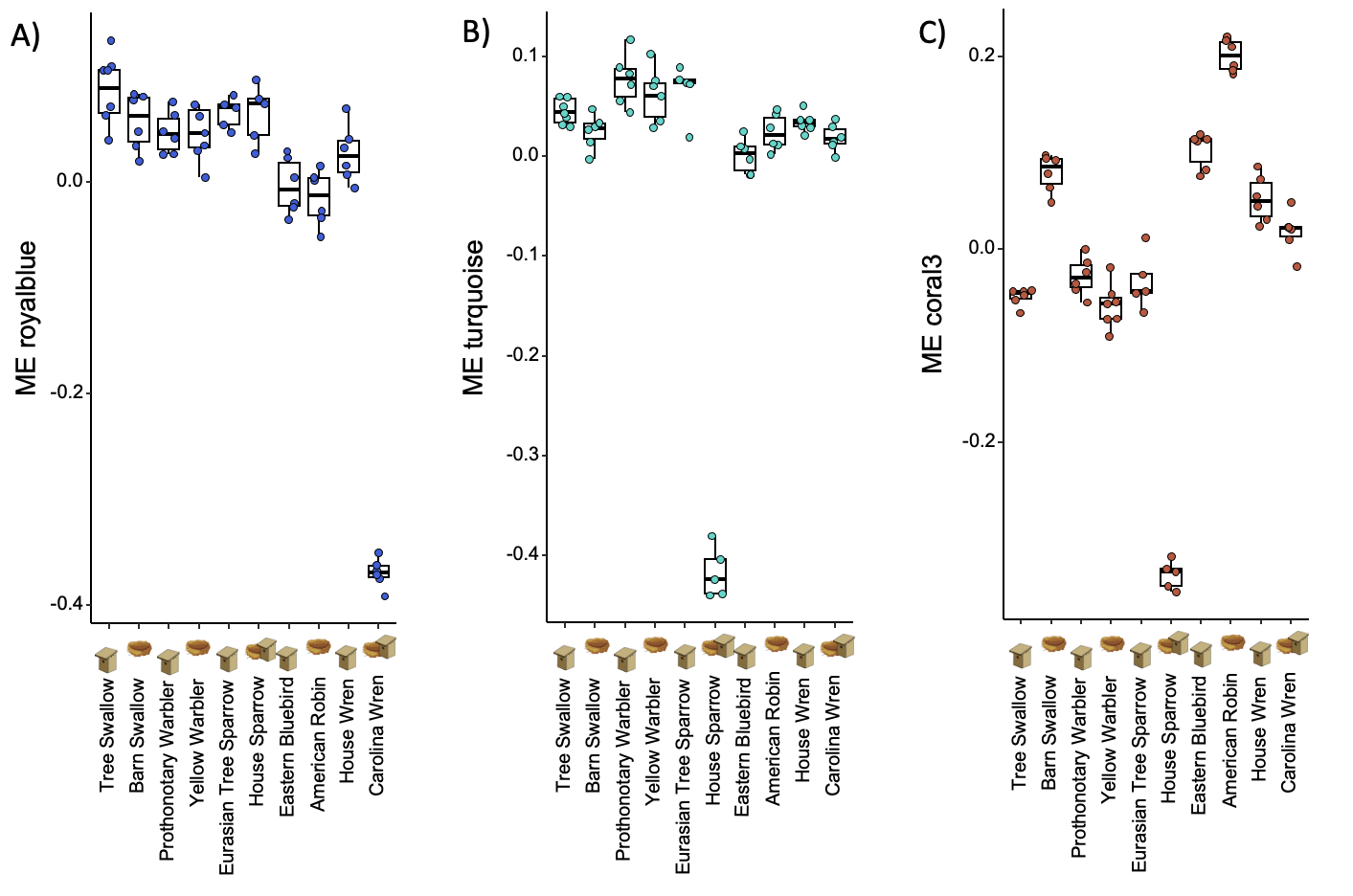

#### **Figure S14.** For females only, WGCNA module eigengenes associated with nest strategy and aggression. A) Gene expression in the royalblue module was significantly associated with the interaction between nest strategy and aggression, for both the 2-strategy and 3-strategy models. This pattern appears driven by Carolina wrens. B) Gene expression in the turquoise module was significantly associated with facultative cavity-nesting, and with the interaction of facultative cavity-nesting and aggression, for the 3-strategy model. This pattern appears driven by house sparrows. C) Gene expression in the coral3 module was significantly associated with facultative cavity-nesting, for the 3-strategy model. This pattern appears driven by house sparrows. Box and whisker plots depict 10th, 25th, 50th, 75th, and 90th percentiles.

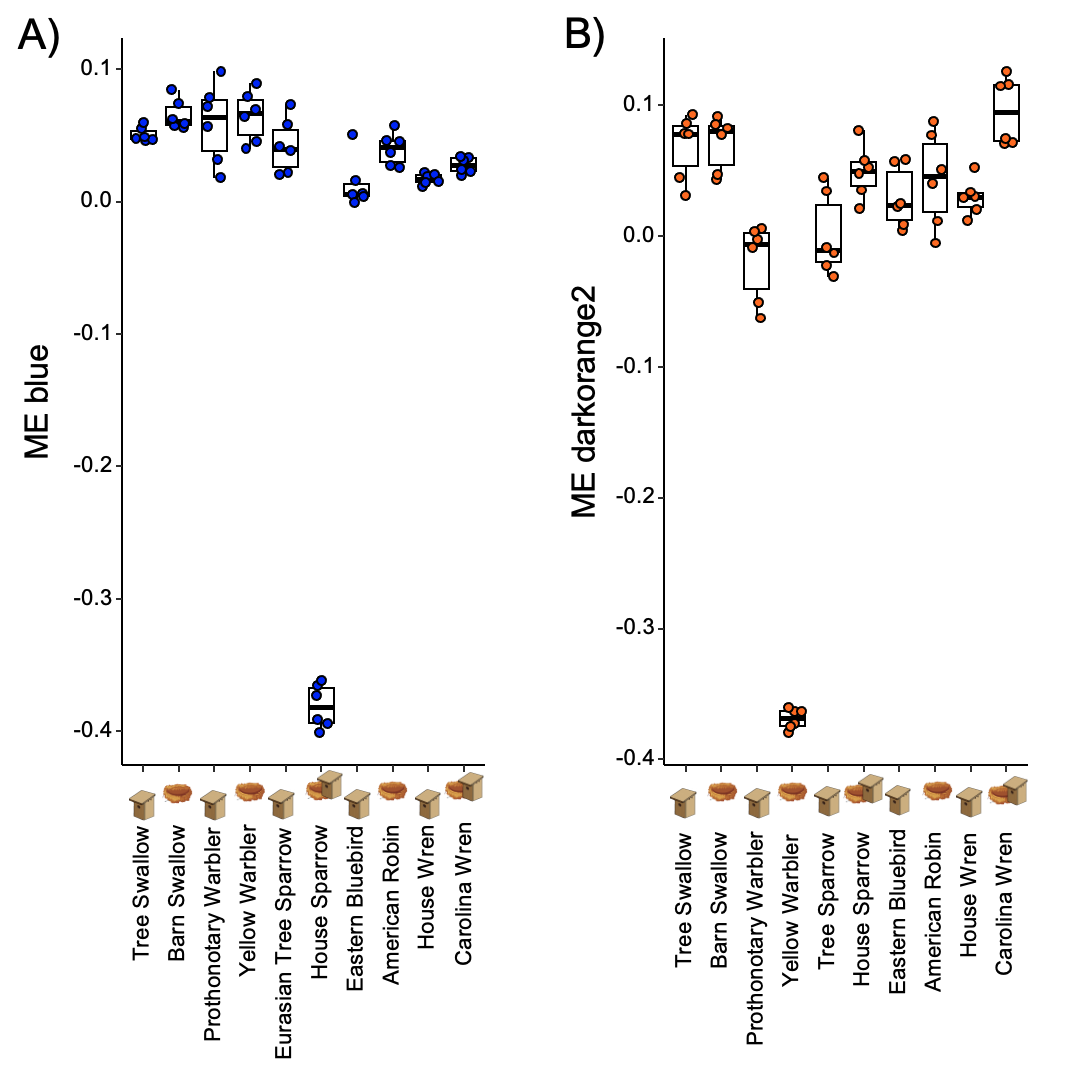

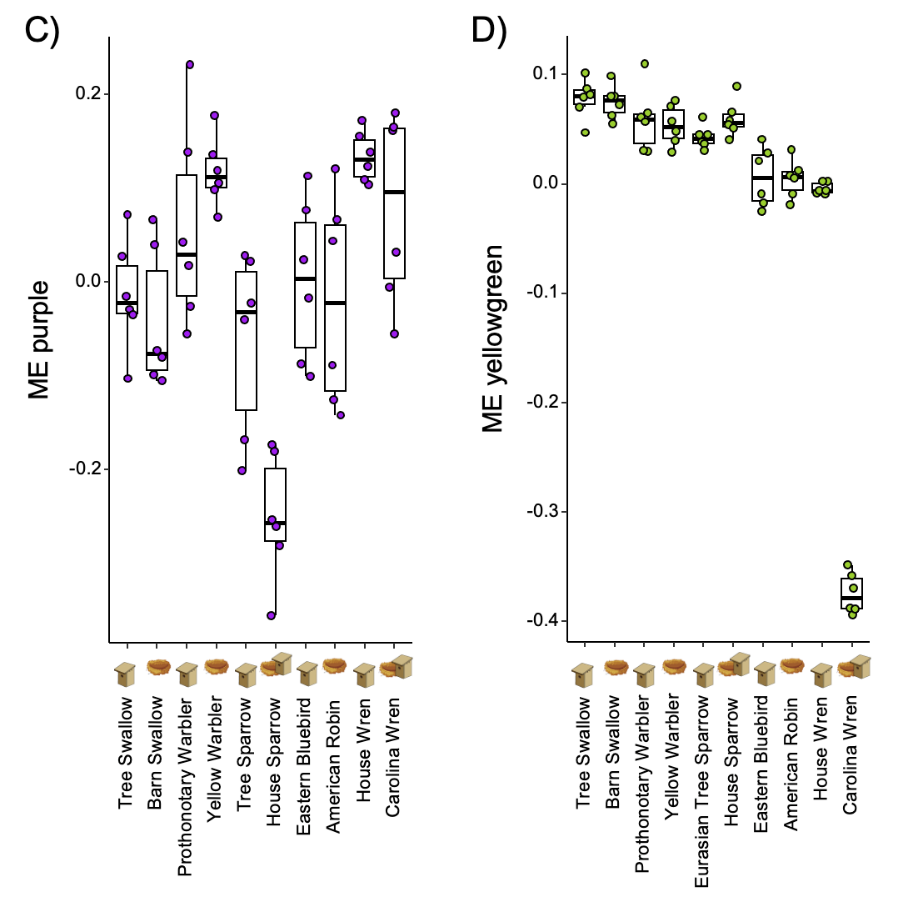

#### **Figure S15.** For males only, WGCNA module eigengenes associated with nest strategy and aggression A) Gene expression in the blue module was significantly associated with nest strategy and with the interaction between nest strategy and aggression, for the 3-strategy model. This pattern appears driven by the house sparrow. B) Gene expression in the darkorange2 module was significantly associated with the interaction between nest strategy and aggression, for 3-strategy model. This pattern appears driven by the yellow warbler. C) Gene expression in the purple module was significantly associated with nest strategy and aggression, for 3-strategy model. This pattern appears driven by the house sparrow. D) Gene expression in the yellowgreen module was significantly associated with the interaction of nest strategy and aggression, for 3-strategy model. This pattern appears driven by the Carolina wren. Box and whisker plots depict 10th, 25th, 50th, 75th, and 90th percentiles.

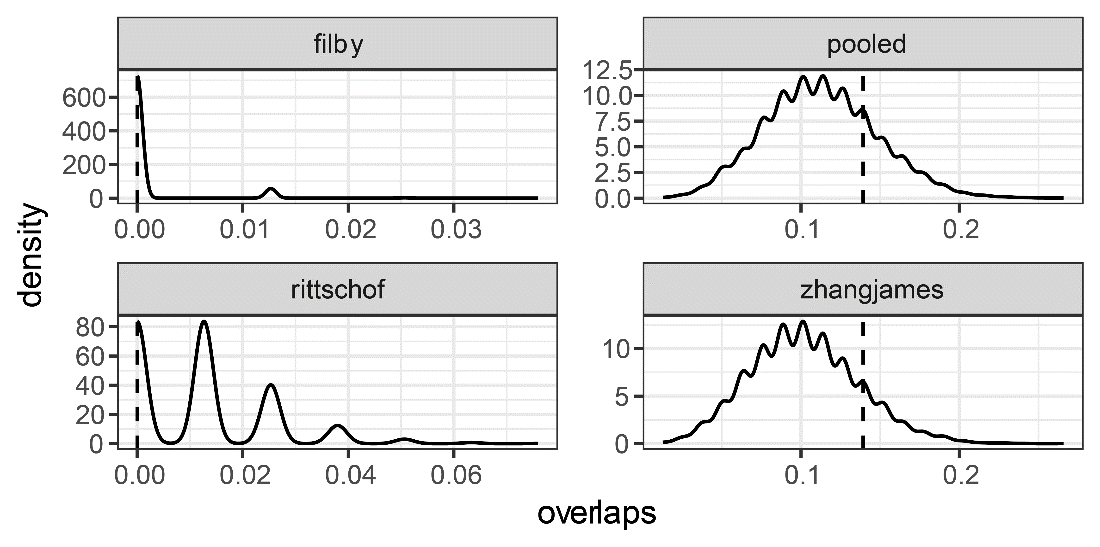

#### **Figure S16.** Randomized overlap distributions between genes significantly associated with aggression (2-level analysis) and candidate gene datasets from Filby (36), Rittschof (34), Zhang-James (35), and all three studies pooled together. X-axis indicates the proportion of overlapping genes between randomized (solid curve) or observed (vertical dashed line) gene sets and the candidate gene dataset (boxes). Stronger than expected overlap with our observed aggression genes is indicated by a dashed line further to the right of the distribution of null values.

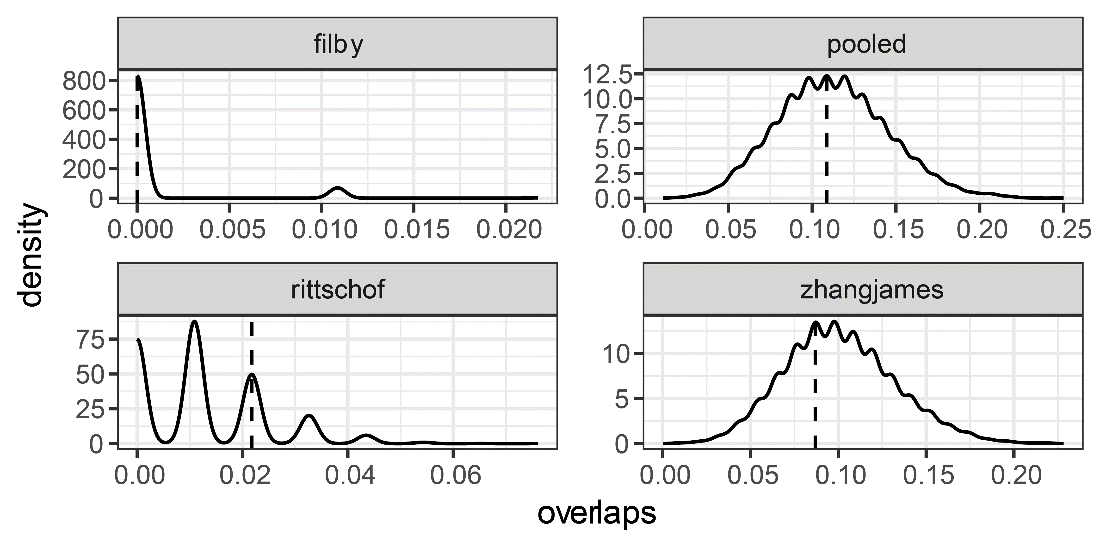

#### **Figure S17.** Randomized overlap distributions between genes significantly associated with aggression (3-level analysis) and candidate gene datasets from Filby (36), Rittschof (34), Zhang-James (35), and all three studies pooled together. X-axis indicates the proportion of overlapping genes between randomized (solid curve) or observed (vertical dashed line) gene sets and the candidate gene dataset (boxes). Stronger than expected overlap with our observed aggression genes is indicated by a dashed line further to the right of the distribution of null values.

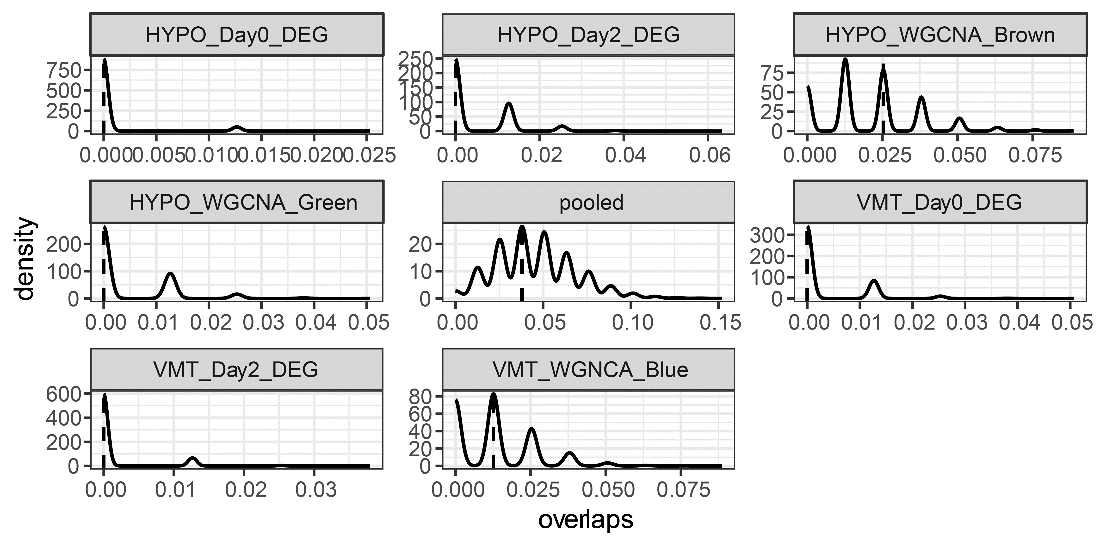

#### **Figure S18.** Randomized overlap distributions for genes significantly associated with aggression (2-level analysis), against aggression candidate gene datasets from Bentz (37). X-axis indicates the proportion of overlapping genes between randomized (solid curve) or observed (vertical dashed line) gene sets and the candidate gene dataset (boxes). Stronger than expected overlap with our observed aggression genes is indicated by a dashed line further to the right of the distribution of null values.

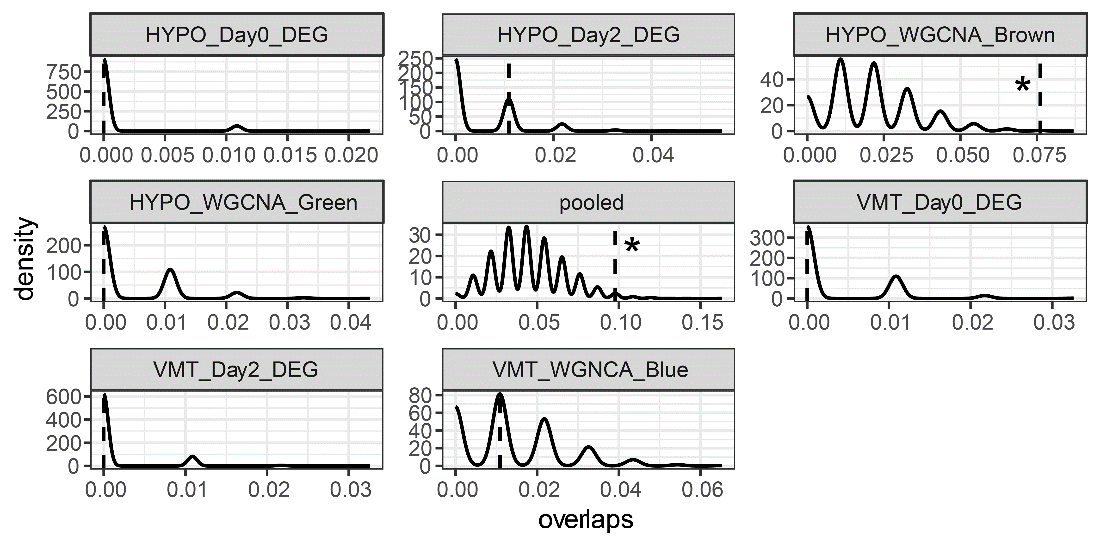

#### **Figure S19.** Randomized overlap distributions for genes significantly associated with aggression (3-level analysis), against aggression-related gene datasets from Bentz (37). X-axis indicates the proportion of overlapping genes between randomized (solid curve) or observed (vertical dashed line) gene sets and the candidate gene dataset (boxes). Stronger than expected overlap with our observed aggression genes is indicated by a dashed line further to the right of the distribution of null values. * indicates significant overlap.
